# Sex-peptide targets distinct higher order processing neurons in the brain to induce the female post-mating response

**DOI:** 10.1101/2024.04.24.590874

**Authors:** Mohanakarthik P. Nallasivan, Deepanshu N.D. Singh, Mohammed Syahir R. S. Saleh, Matthias Soller

## Abstract

Sex-peptide (SP) transferred during mating induces female post-mating responses including refractoriness to re-mate and increased oviposition in *Drosophila*. Yet, where SP-target neurons reside, remained uncertain. Here we show that expression of membrane-tethered SP (mSP) pre-dominantly in the head or trunk either reduces receptivity or increases oviposition, respectively. Using fragments from large regulatory regions of *Sex Peptide Receptor*, *fruitless* and *doublesex* genes together with intersectional expression of mSP, we identified distinct interneurons in the brain and abdominal ganglion controlling receptivity and oviposition. These SP Response Inducing Neurons (SPRINz) can induce post-mating responses through SP received by mating. Trans-synaptic mapping of neuronal connections reveals input from sensory processing neurons and two post-synaptic trajectories as output. Hence, SP-target neurons operate as key integrators of sensory information for decision of behavioural outputs. Multi-modularity of SP-targets further allows females to adjust SP-mediated male manipulation to physiological state and environmental conditions for maximizing reproductive success.

## Introduction

Reproductive behaviors are to a large degree hard-wired in the brain to guarantee reproductive success making the underlying neuronal circuits amenable to genetic analysis (Dulac and Kimchi 2007; Yamamoto and Koganezawa 2013; Anderson 2016; Rings and Goodwin 2019).

During development, sex-specific circuits are built into the brain under the control of the sex determination genes *doublesex* (*dsx*) and *fruitless* (*fru*) in *Drosophila* (Schutt and Nothiger 2000; Billeter et al. 2006). They encode transcription factors that are alternatively spliced in a male or female specific mode (Schutt and Nothiger 2000). By default, the *dsx* gene generates the male-specific isoform Dsx^M^, while a female-specific isoform Dsx^F^ is generated by alternative splicing and expressed in about ∼700 distinct neurons in the brain important for female reproductive behaviors directing readiness to mate and egg laying (Rideout et al. 2010; Rezaval et al. 2012). Fru^M^ is expressed in about ∼1000 neurons in males and implements development of neuronal circuitry key to display male courtship behavior, but is switched off in females through alternative splicing by incorporation of a premature stop codon (Demir and Dickson 2005; Manoli et al. 2005; Stockinger et al. 2005).

The circuitry of female specific behaviors including receptivity to courting males for mating and egg laying have been mapped using intersectional gene expression via the *split-GAL4* system to restrict expression of activators or inhibitors of neuronal activity to very few neurons (Aranha and Vasconcelos 2018; Wang et al. 2020a; Wang et al. 2020b; Wang et al. 2021; Cury and Axel 2023). Through this approach, sensory neurons in the genital tract have been identified as key signal transducers for the readiness to mate and the inhibition of egg laying connecting to central parts of the brain via projection to abdominal ganglion neurons (Hasemeyer et al. 2009; Yang et al. 2009; Rezaval et al. 2012; Feng et al. 2014). This circuit then projects onto centrally localized pattern generators in the brain to direct a behavioral response via efferent neurons (Wang et al. 2020a; Wang et al. 2020b; Wang et al. 2021).

Once females have mated, they will reject courting males and lay eggs (Manning 1967). Post-mating responses (PMRs) are induced by male derived sex-peptide (SP) and other substances transferred during mating (Chen et al. 1988; Avila et al. 2011; Hopkins and Perry 2022; Kim et al. 2024; Singh and Soller 2025). In addition to refractoriness to remate and oviposition, SP will induce a number of other behavioral and physiological changes including increased egg production, feeding, a change in food choice, sleep, memory, constipation, midgut morphology, stimulation of the immune system, and sperm storage and release (Soller et al. 1999; Peng et al. 2005; Carvalho et al. 2006; Domanitskaya et al. 2007; Kim et al. 2010; Ribeiro and Dickson 2010; Scheunemann et al. 2019) (Cognigni et al. 2011) (Avila et al. 2010; Isaac et al. 2010; Wainwright et al. 2021; White et al. 2021). SP binds to broadly expressed Sex Peptide Receptor (SPR), an ancestral receptor for myoinhibitory peptides (MIPs) (Yapici et al. 2008; Kim et al. 2010; Jang et al. 2017). Although MIPs seem not to induce PMRs, excitatory activity of MIP expressing neurons underlies re-mating (Yapici et al. 2008; Kim et al. 2010; Jang et al. 2017). Expression of membrane-tethered SP (mSP) induces PMRs in an autocrine fashion when expressed in neurons, but not glia (Nakayama et al. 1997; Haussmann et al. 2013).

First attempts to identify SP target neurons by enhancer *GAL4* induced expression of *UASmSP* only identified lines with broad expression in the nervous system (Nakayama et al. 1997). Later, drivers with more restricted expression including *dsx*, *fru* and *pickpocket* (*ppk*) genes were identified, but they are expressed in all parts of the nervous system throughout the body eluding to reveal the location of SP target sites unambiguously (Yapici et al. 2008; Hasemeyer et al. 2009; Yang et al. 2009; Rezaval et al. 2012; Haussmann et al. 2013).

To delineate where in the *Drosophila* SP target neurons are located which induce the main PMRs, refusal to mate and egg laying, we expressed mSP pre-dominantly in the head or trunk. These experiments separate reduction of receptivity induced in the head from trunk induction of egg laying. To further restrict our search for SP target neurons, we focused on three genes, *SPR*, *dsx* and *fru,* because SPR is broadly expressed but anticipated to induce PMRs only from few neurons, and because *GAL4* inserted in the endogenous *dsx* and *fru* loci induces PMRs from mSP expression. Using *GAL4* tiling lines with fragments encompassing the regulatory regions of complex *SPR*, *fru* and *dsx* genes (Pfeiffer et al. 2008; Jenett et al. 2012; Kvon et al. 2014), we identified one regulatory region in each gene reducing receptivity and inducing egg laying upon mSP expression, and one additional region in *SPR* only inducing egg laying. To further refine this analysis, we used intersectional gene expression using *split-GAL4* and *flipase* (*flp*) mediated excision of stop cassettes in *UAS* reporters (Struhl and Basler 1993; Luan et al. 2006). Consistent with previous results that the SP response can be induced via multiple pathways (Haussmann et al. 2013), we found distinct sets of SP Response Inducing Neurons (SPRINz) in the central brain and the abdominal ganglion that can induce PMRs via expression of mSP either reducing receptivity and inducing egg laying, or affecting only one of these PMRs. In contrast, we identified genital tract neuron expressing lines including *splitGAL4 nSyb* ∩ *ppk* that did not induce PMRs by expression of mSP. Likewise, we find expression of mSP or neuronal activation in head Sex Peptide Sensing Neurons (SPSN) neurons can induce PMRs. Mapping the pre- and post-synaptic connections of the distinct SP target neurons by *retro-* and *trans*-Tango (Talay et al. 2017; Sorkaç et al. 2023) revealed that SP target neurons direct higher order sensory processing in the central brain. These neurons feed into two common post-synaptic neuronal subtypes indicating that SP interferes with the integration of diverse sensory inputs to build a stereotyped output either reducing receptivity and/or increasing egg laying.

## Results

### Reduction of receptivity and induction of egg laying are separable by head and trunk expression of membrane-tethered SP

Due to the complex behavioral and physiological changes induced by SP, neurons in the central nervous system have been suspected as main targets for SP (Kubli 1992). To express mSP only in the head we used an *elav FRTstopFRT GAL4* in combination with *otdflp*, that expresses in the head to drive recombination and head-specific expression of mSP from *UAS* (Figure 1A, Figures S1A-F) (Haussmann et al. 2008; Asahina et al. 2014; Zaharieva et al. 2015; Nallasivan et al. 2021). To express mSP predominantly in the trunk we used *tshGAL4* (Figure 1B, Figures S1G-L) (Soller et al. 2006).

**Figure 1:**
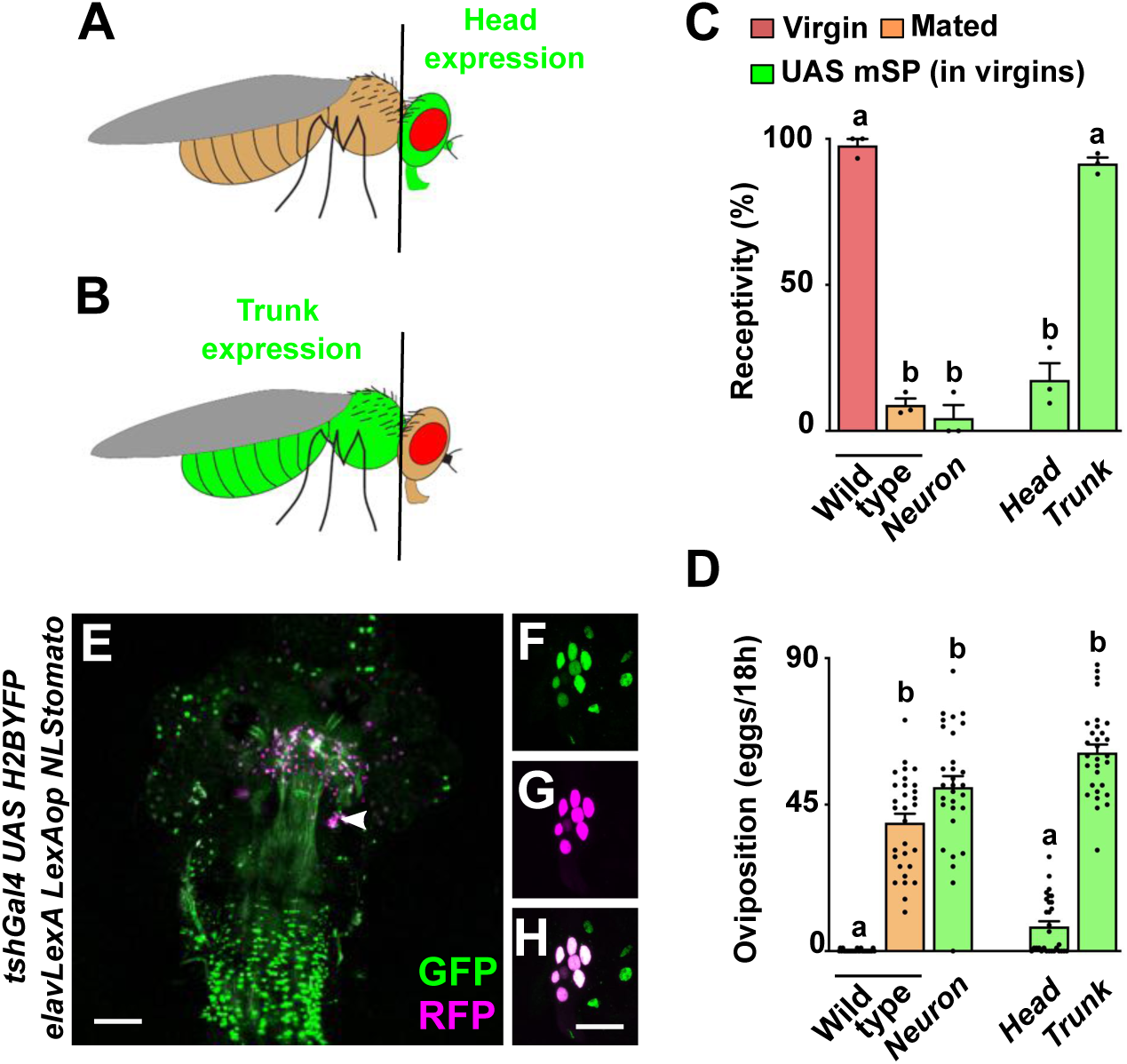
The main PMRs in females can be separated. **A, B)** Schematic depiction of head and trunk expression in *Drosophila elav FRTstopFRT GAL4*; *otdflp* (A) and in *tshGAL4* (B) visualized by *UAS GFP* (green). **C, D)** Receptivity (C) and oviposition (D) of wild type control virgin (red) and mated (orange) females, and virgin females expressing *UAS mSP* (green) pan-neuronally with *nsybGAL4* or in head and trunk patterns shown as means with standard error from three repeats for receptivity (21 females per repeat) by counting the number of females mating within a 1 h period or for oviposition by counting the eggs laid within 18 hours from 30 females. Statistically significant differences from ANOVA post-hoc comparison are indicated by different letters (p<0.0001). **E-H)** Representative adult female genital tract showing *tshGAL4 UAS H2BYFP* (green) and *elavLexA LexAop NLStomato* (red) nuclear expression. The magnification (F-H) shows sensory genital tract neurons. Scale bar shown in E and H are 100 μm and 20 μm, respectively.

When we expressed mSP in the head, females reduced receptivity indistinguishable from mated females, but did not lay eggs, thereby again demonstrating that the two main PMRs can be separated (Figures 1C and 1D) (Haussmann et al. 2013). In contrast, when we expressed mSP in the trunk, females remained receptive, but laid eggs in numbers indistinguishable from mated females (Figures 1C and 1D).

Moreover, *tshGAL4* is expressed in *fru*, *dsx*, *ppk* genital tract sensory neurons (Figures 1E-1H). Since mSP expression with *tshGAL4* does not affect receptivity, these genital tract neurons unlikely are direct targets for SP (Haussmann et al. 2013). Taken together, these results indicate presence of SP target neurons in the brain and ventral nerve cord (VNC) for the reduction of receptivity and induction of egg laying, respectively.

### Few restricted regulatory regions in large *SPR*, *fru* and *dsx* genes can induce the SP response

Expression of mSP from *UAS* via *GAL4* inserts in *fru* and *dsx* genes induces a robust reduction in receptivity and increase in egg laying (Rezaval et al. 2012; Haussmann et al. 2013). To identify SP target neurons, we thought to dissect the broad expression pattern of complex *SPR*, *fru* and *dsx* genes spanning 50-80 kb by identifying regulatory DNA fragments in the enhancer regions that drive *UAS mSP* in a subset of neurons. For these experiments, we analysed 22, 27 and 25 *GAL4* lines from the VDRC and Janelia tiling *GAL4* projects (Pfeiffer et al. 2008; Jenett et al. 2012; Kvon et al. 2014) (Figures 2A-2C).

**Figure 2:**
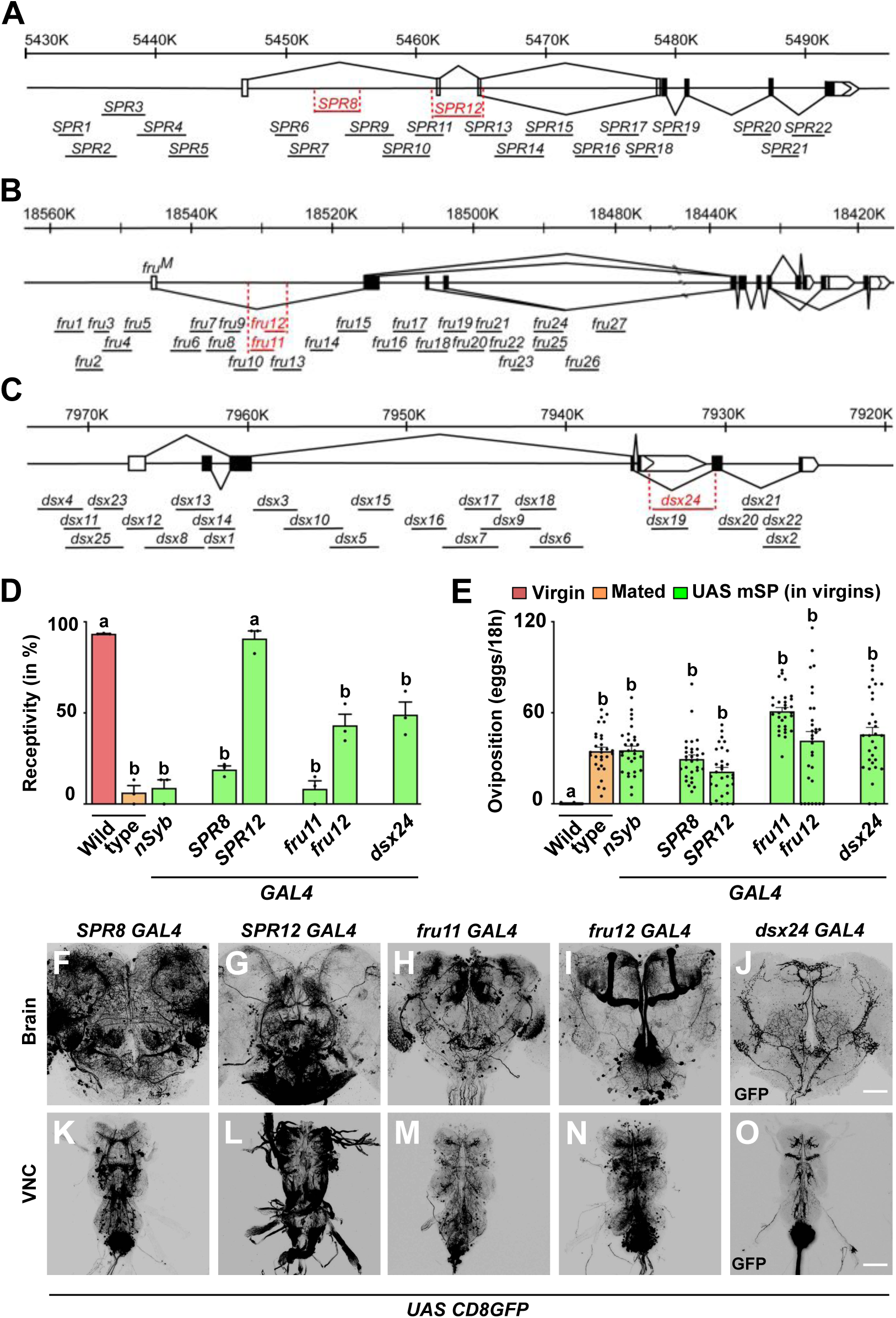
Distinct regulatory regions in *SPR*, *fru* and *dsx* genes induce PMRs from mSP expression. **A-C)** Schematic representation of *SPR*, *fru*, and *dsx* chromosomal regions depicting coding and non-coding exons as black or white boxes, respectively, and splicing patterns in solid lines. Vertical lines below the gene model depict enhancer *GAL4* lines with names and those in red showed PMRs by expression of mSP. **D, E)** Receptivity (D) and oviposition (E) of wild type control virgin (red) and mated (orange) females, and virgin females expressing *UAS mSP* (green) under the control of GAL4 pan-neuronally in *nsyb* or in *SPR8*, *SPR12*, *fru11, fru12*, and *dsx24* patterns shown as means with standard error from three repeats for receptivity (21 females per repeat) by counting the number of females mating within a 1 h period or for oviposition by counting the eggs laid within 18 hours from 30 females. Statistically significant differences from ANOVA post-hoc comparison are indicated by different letters (p≤0.0001). **F-O)** Representative adult female brains (F-J) and ventral nerve cords (VNC, K-O) expressing *UAS CD8GFP* under the control of *SPR8*, *SPR12*, *fru11*, *fru12* and *dsx24 GAL4*. Scale bars shown in J and O are 50 µm and 100 µm, respectively.

Strikingly, in *SPR*, *fru* and *dsx* genes we identified only one regulatory region in each gene (*SPR8*, *fru11/12* and *dsx24*) that reduced receptivity and induced egg laying through *GAL4 UAS* expression of mSP (Figures 2D and 2E). In addition, we identified one line (*SPR12*) in the *SPR* gene, that induced egg laying, but did not reduce receptivity consistent with previous results that SP regulation of receptivity and egg laying can be split (Haussmann et al. 2013).

All of these lines expressed in subsets of neurons in the central brain and the ventral nerve cord in distinct, but reduced patterns compared to the expression of the *SPR*, *fru* and *dsx* genes (Yapici et al. 2008; Rideout et al. 2010; Zhou et al. 2014) (Figures 2F-2O). Moreover, these lines showed prominent labelling of abdominal ganglion neurons in the VNC (Figures 2K-2O). In addition, all of these lines except *SPR12* are also expressed in genital tract sensory neurons (Figure S2).

From all the 74 lines that we have analyzed for PMRs from *SPR*, *fru* and *dsx* genes, we also analysed expression in genital tract sensory neurons as they had been postulated to be the primary targets of SP (Yapici et al. 2008; Hasemeyer et al. 2009; Yang et al. 2009; Rezaval et al. 2012). Apart from PMR inducing lines *SPR8*, *fru11*, *fru12* and *dsx24*, that showed expression in genital tract sensory neurons, we identified three lines (*SPR3*, *SPR 21* and *fru9*), which also robustly expressed in genital tract sensory neurons but did not induce PMRs from expression of mSP (Figure S3, S4E, S4F, S4I and S4J).

### Secondary ascending abdominal ganglion neurons can induce the PMRs from mSP expression

A screen aiming to identify neurons involved in the control of receptivity and egg laying by expression of the rectifying potassium channel Kir2.1 identified six enhancer *GAL4* driver lines (*FD1-6*) (Feng et al. 2014). *FD1-6* are expressed in diverse subsets of neurons in the brain and the ventral nerve cord, in particular they show common expression in the abdominal ganglion with projections to the central brain. The lines expressing in FD1-5 neurons have been termed SAG (secondary ascending abdominal ganglion neurons) neurons, that are also interconnected with myoinhibitory peptide sensing neurons (Jang et al. 2017). Since enhancer lines identified in *SPR*, *fru* and *dsx* genes are prominently expressed in the abdominal ganglion, we tested whether mSP expression from these FD1-6 lines induced PMRs.

From these six lines, one robustly suppressed receptivity and induced egg laying (*FD6/VT003280*), while two lines only induced egg laying (*FD3/VT4515* and *FD4/V000454*) similar to controls from mSP expression (Figure 3). Again, all three lines also expressed in subsets of neurons in the central brain and VNC, particularly in the abdominal ganglion (Zhou et al. 2014). In addition, *FD3* and *FD4* did not express in genital tract sensory neurons, in contrast to *FD6* (Feng et al. 2014). A *SAG1 split-GAL4* (*VT050405/FD1 AD* and *VT007068/FD2 DBD*) did not show a response to expression of mSP and virgin females, e.g. they mated and did not lay eggs (Figure 3).

**Figure 3:**
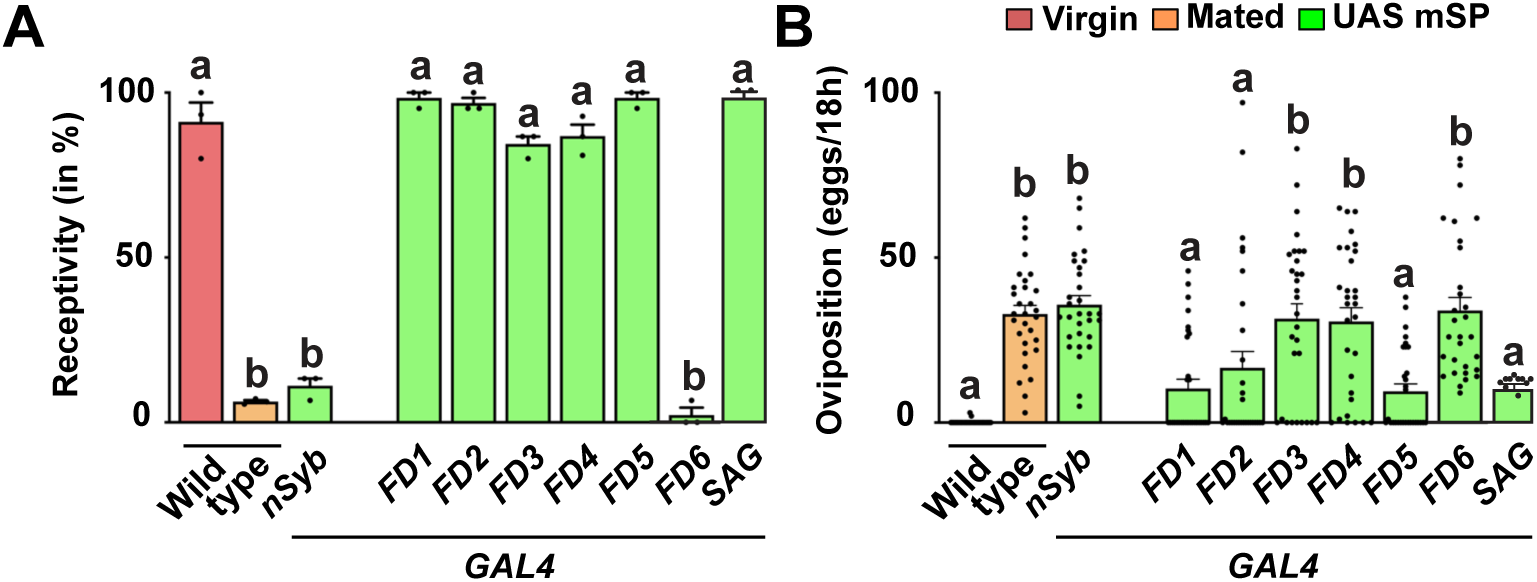
Expression of mSP in secondary ascending abdominal ganglion neurons induces PMRs. **A, B)** Receptivity (A) and oviposition (B) of wild type control virgin (red) and mated (orange) females, and virgin females expressing *UAS mSP* (green) under the control of GAL4 pan-neuronally in *nsyb* or in *FD1*, *FD2*, *FD3*, *FD4*, *FD5*, and *FD6,* or with *SAG split-Gal4* patterns shown as means with standard error from three repeats for receptivity (21 females per repeat) by counting the number of females mating within a 1 h period or for oviposition by counting the eggs laid within 18 hours from 30 females. Statistically significant differences from ANOVA post-hoc comparison are indicated by different letters (p<0.0001).

### Intersectional expression reveals distinct mSP responsive neurons in the central brain and abdominal ganglion

To further restrict the expression to fewer neurons, we intersected the expression patterns of those lines that induced robust reduction of receptivity and increase of egg laying using *split-GAL4* (*SPR8*, *fru11/12*, *dsx* and *FD6*, for further experiments we used *dsxGAL4-DBD*, because *dsx24* is less robust and *fru11* and *fru12* were made into one fragment) that activates the *UAS* reporter when *GAL4* is reconstituted via dimerization of activation (AD-GAL4) and DNA binding (GAL4-DBD) domains (Luan et al. 2006) (Figure 4A).

**Figure 4:**
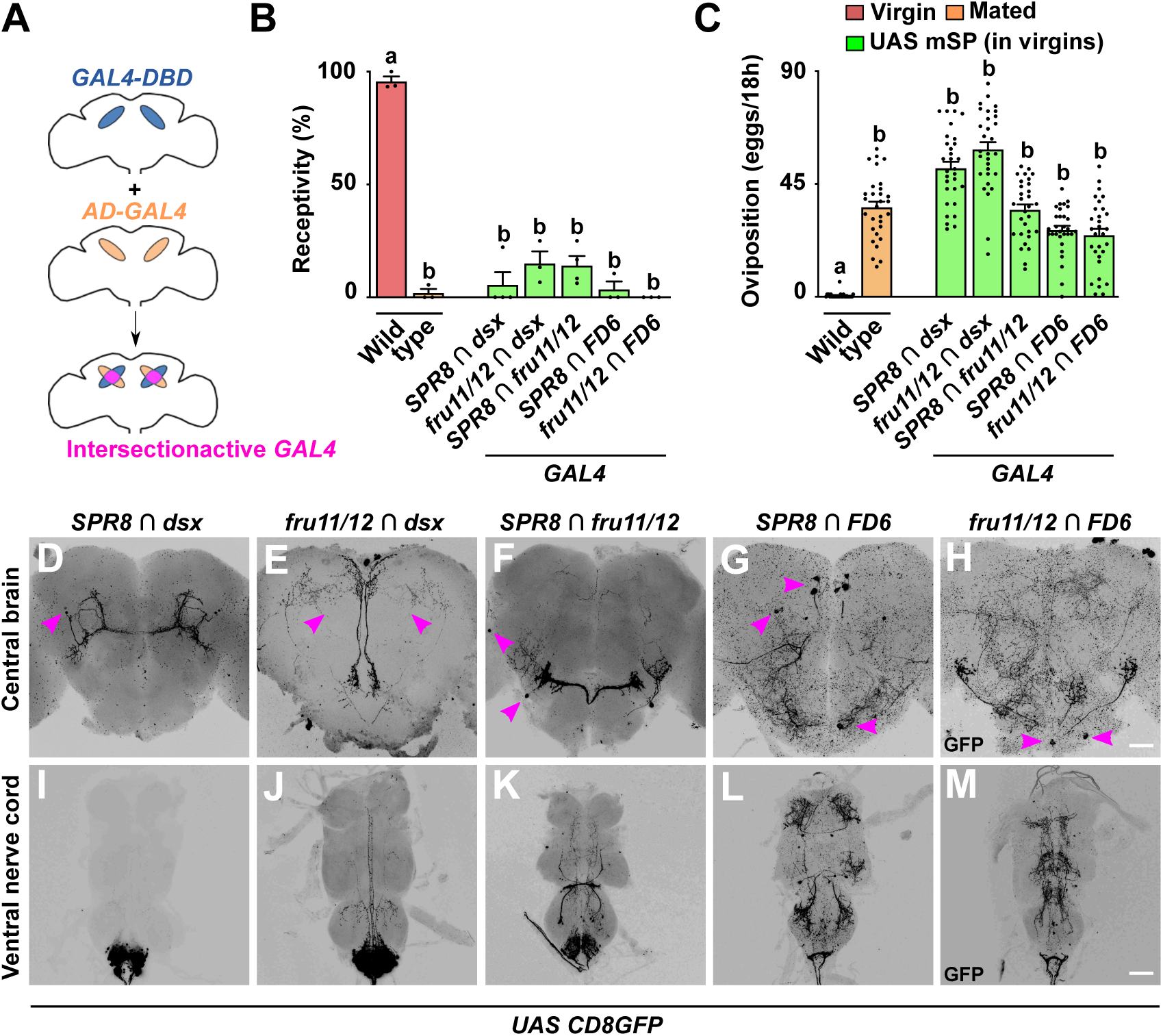
Distinct circuits from intersection of *SPR*, *fru*, *dsx* and *FD6* patterns in the brain and VNC induce PMRs from mSP expression. **A)** Schematic showing the intersectional gene expression approach: GAL4 activation (AD, orange) and DNA binding domains (DBD, blue) are expressed in different, but overlapping patterns. Leucine zipper dimerization reconstitutes a functional split-GAL4 in the intersection (pink) to express *UAS* reporters. **B, C)** Receptivity (B) and oviposition (C) of wild type control virgin (red) and mated (orange) females, and virgin females expressing *UAS mSP* (green) under the control of *split-GAL4* intersecting *SPR8* ∩ *fru11/12, SPR8* ∩ *dsx, SPR8* ∩ *FD6, fru11/12* ∩ *dsx* and *fru11/12* ∩ *FD6* patterns shown as means with standard error from three repeats for receptivity (21 females per repeat) by counting the number of females mating within a 1 h period or for oviposition by counting the eggs laid within 18 hours from 30 females. Statistically significant differences from ANOVA post-hoc comparison are indicated by different letters (p<0.0001). **D-M)** Representative adult female brains and ventral nerve cords (VNC) expressing *UAS CD8GFP* under the control of *SPR8* ∩ *fru11/12, SPR8* ∩ *dsx, SPR8* ∩ *FD6, fru11/12* ∩ *dsx* and *fru11/12* ∩ *FD6*. Scale bars shown in H and M are 50 µm and 100 µm, respectively.

Again, intersection of *SPR8* with *fru11/12*, *dsx* or *FD6*, and *fru11/12* with *dsx* or *FD6* expression robustly reduced receptivity and increased egg laying upon expression of mSP (Figures 4B and 4C). Accordingly, we termed these neurons SP Response Inducing Neurons (SPRINz), though the exact identity in the *splitGAL4* intersection population needs to be determined.

When we further analyzed the expression of these *split-GAL4* intersections in the brain, we found that each combination first showed very restricted expression, but second, that none of these combinations labeled the same neurons (Figures 4D-4H). For *dsx* neurons, *split-GAL4* intersections correspond to a subset of dPC2l (*SPR8* ∩ *dsx*) and dPCd-2 (*fru11/12*∩ *dsx*) neurons (Deutsch et al. 2020; Schretter et al. 2020; Nojima et al. 2021). These results suggest the SP targets interneurons in the brain that feed into higher processing centers from different entry points likely representing different sensory input.

In the ventral nerve cord, we found expression in the abdominal ganglion with all *split-GAL4* combinations (Figures 4I-4M). In particular, intersection of *dsx* with *SPR8* or *fru11/12* showed exclusive expression in the abdominal ganglion, while the other combinations also expressed in other cells of the VNC. All together, these data suggest that the abdominal ganglion harbors several distinct type of neurons involved in directing PMRs (Oliveira-Ferreira et al. 2023).

In the female genital tract, these *split-Gal4* combinations show expression in genital tract neurons with innervations running along oviduct and uterine walls (Figures S5A-S5E). In addition, *SPR8* lZ fru11/*12* and *SPR8* lZ *dsx* were also expressed in the spermathecae (Figures S5A-S5B).

### mSP responsive neurons rely on SPR and are required for PMRs induced by SP delivered through mating

Next, we tested whether PMRs induced by mSP expression in the *SPR8* ∩ *dsx, fru11/12* ∩ *dsx* or *SPR8* ∩ *fru11/12* rely on *SPR*. Expression of mSP in *dsx* ∩ *SPR8* and *dsx* ∩ *fru11/12* neurons in *SPR* mutant females did not reduce receptivity or induce egg laying (Figures 5A and B, see also Figures 4A and B), while a partial response was observed for *SPR8* ∩ *fru 11/12* induced mSP expression in *SPR* mutant females, which is consistent with presence of additional receptors for SP (Haussmann et al. 2013).

**Figure 5:**
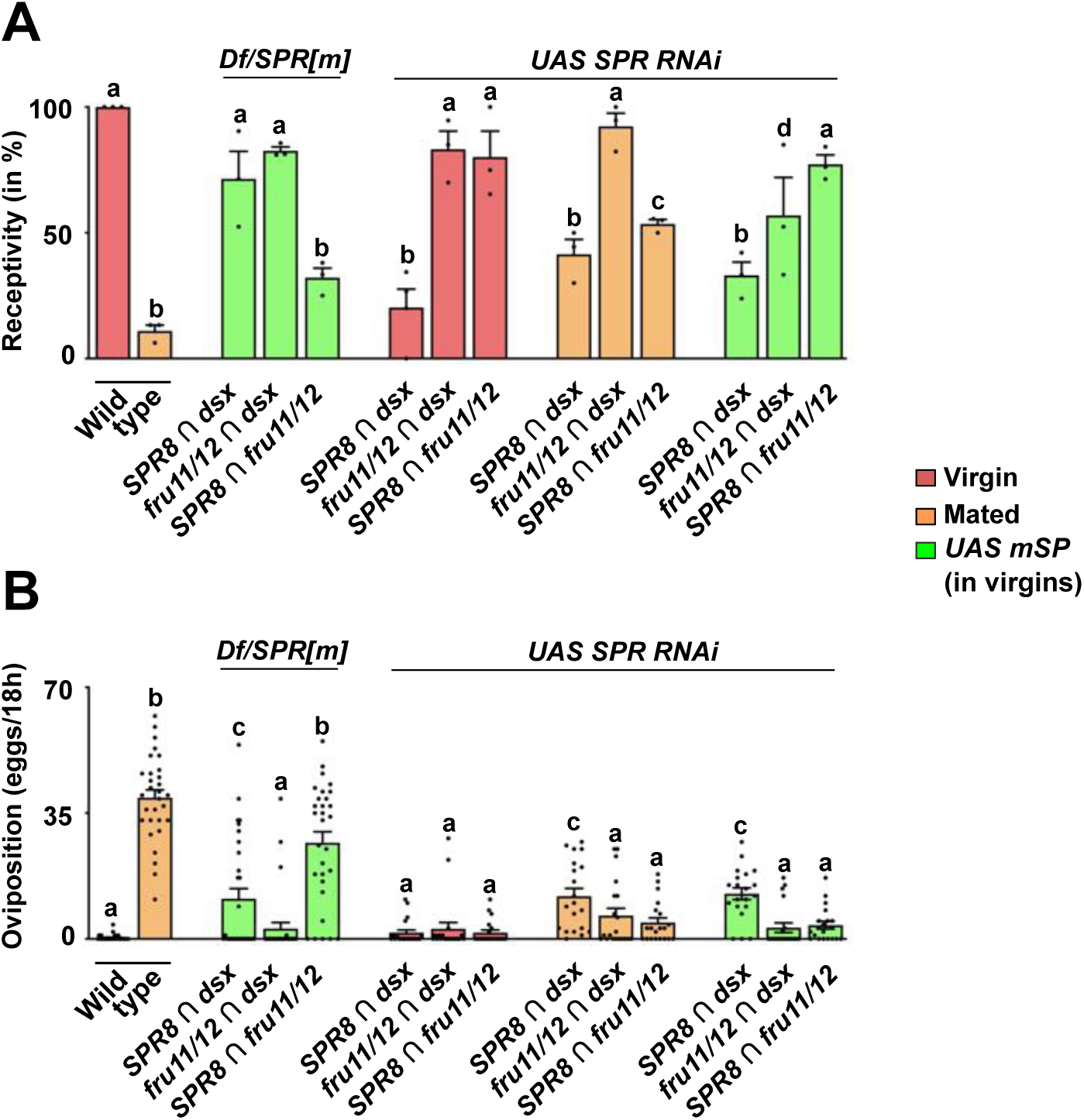
Distinct neuronal circuitries from intersection of *SPR*, *fru* and *dsx* sense SP after mating to induce PMRs. **A, B)** Receptivity (A) and oviposition (B) of wild type control virgin (red) and mated (orange) females, and virgin females expressing *UAS mSP* (green) under the control of *split-Gal4* intersecting *SPR8* ∩ *dsx, fru11/12* ∩ *dsx,* and *SPR8* ∩ *fru11/12* patterns in *SPR/Df* mutant females or *SPR* RNAi knock-down shown as means with standard error from three repeats for receptivity (21 females per repeat) by counting the number of females mating within a 1 h period or for oviposition by counting the eggs laid within 18 hours from 30 females. Statistically significant differences from ANOVA post-hoc comparison are indicated by different letters (p<0.0001 except p=0.002 and p=0.006 for c and d in A, and p=0.004 for c in B).

Since SP is transferred during mating to females and enters the hemolymph (Haussmann et al. 2013), we wanted to test whether SPR is required in these neurons for inducing PMRs after mating. For *SPR RNAi* in *dsx* ∩ *fru11/12* and *SPR8* ∩ *fru 11/12* neurons, no reduction, or a partial reduction, of receptivity was observed, respectively, while *SPR RNAi* in *dsx* ∩ *SPR8* neurons turned virgin females unreceptive (Figure 5A). Expression of mSP in *dsx* ∩ *fru11/12* neurons in the context of *SPR RNAi* partially reduced receptivity again suggesting additional receptors for SP (Haussmann et al. 2013).

Strikingly, however, *SPR RNAi* in these neurons prevented egg laying independent of whether SP was delivered by mating or when tethered to the membrane of these neurons (Figure 5B).

These results demonstrate that neurons identified by *split-GAL4* intersected expression of *SPR8* with *dsx* or *fru11/12*, or *fru11/12* with *dsx* are genuine SP targets as they rely on *SPR* and PMRs are induced by SP delivered through mating.

### Expression of mSP in distinct neurons in the brain induces PMRs

The analysis of *ppkGAL4* neurons in SP-insensitive *Nup54* alleles revealed a hierarchy of trunk neurons that dominate over central brain neurons (Nallasivan et al. 2021). To focus on the role of central brain neurons, we generated a *UAS mSP* line with a stop cassette (*UAS FRTstopFRT mSP*) that allows to restrict expression of mSP to the head in the presence of *otdflp*, which only expresses in the head (Figure 6A), but not in the trunk (Asahina et al. 2014; Nallasivan et al. 2021).

**Figure 6:**
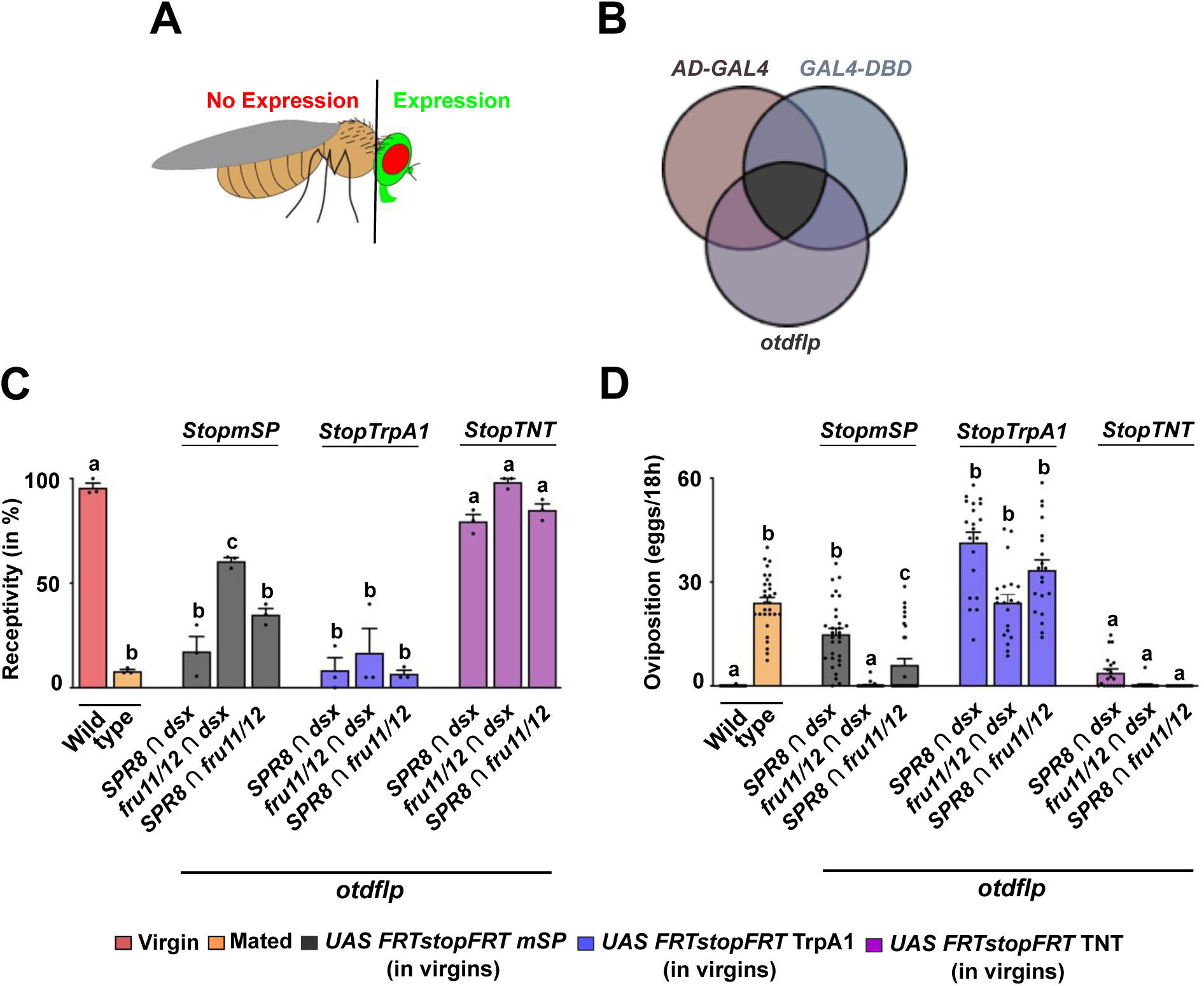
Distinct neuronal circuitries in the brain sense SP to induce PMRs. **A, B)** Schematic depiction of *UAS GFP* (green) expression in the head of *Drosophila* (A) combining *split-GAL4* intersectional expression (*AD-GAL4* and *GAL4-DBD*) with brain-expressed *otdflp* mediated recombination of *UAS FRTGFPstopFRTmSP* (B). **C, D)** Receptivity (C) and oviposition (D) of wild type control virgin (red) and mated (orange) females, and virgin females expressing *UAS FRTGFPstopFRTmSP* (grey), *UAS FRTGFPstopFRTTrpA1* (purple) and *UAS FRTGFPstopFRTTNT* (pink) under the control of *split-GAL4* intersecting *SPR8* ∩ *dsx, fru11/12* ∩ *dsx* and *SPR8* ∩ *fru11/12* patterns with brain-specific FRT-mediated recombination by *otdflp* shown as means with standard error from three repeats for receptivity (21 females per repeat) by counting the number of females mating within a 1 h period or for oviposition by counting the eggs laid within 18 hours from 30 females. Statistically significant differences from ANOVA post-hoc comparison are indicated by different letters (p<0.0001 except p<0.0004 for c in C, p<0.007 for c in D).

In combination with the intersectional approach, we now can restrict mSP expression to few central brain neurons, or alternatively activate or silence these neurons (Figure 6B). Expression of mSP in *SPR8* ∩ *dsx, fru11/12* ∩ *dsx* or *SPR8* ∩ *fru11/12* neurons in the central brain significantly reduced receptivity, but oviposition was only substantially induced in *SPR8* ∩ *dsx* brain neurons (Figures 6C and 6D). In *fru11/12* ∩ *dsx* or *SPR8* ∩ *fru11/12,* PMR inducing neurons from the VNC could be required to potentiate the response.

These results clearly demonstrate a role for brain neurons in the SP response. However, we noticed that the flipase approach can result in false negatives as *fruflp* inserted in the same position in the endogenous locus as *fruGAL4* does not induce a response with *UAS FRTstopFRT* mSP in contrast to *fruGAL4* induced expression of mSP. In contrast, the same experiment with *dsxGAL4* and *dsxflp* results in a positive SP response indistinguishable from mated females (Haussmann et al. 2013).

Next, we tested whether neuronal activation or inhibition would induce a post-mating response. Strikingly, conditional activation of *SPR8* ∩ *dsx, fru11/12* ∩ *dsx* or *SPR8* ∩ *fru11/12* brain neurons with TrpA1in adult females completely inhibited receptivity and induced egg laying comparable to mated females (Figures 6C and 6D). In contrast, inhibition of these neurons with tetanus toxin (TNT) did not alter the virgin state, e.g. receptivity was not reduced and egg laying was not induced (Figures 6C and 6D).

### Genital tract neurons do not mediated changes in receptivity and oviposition by mSP

Since genital tract sensory neurons have been postulated to induce the SP response, we tested previously identified *split-GAL4* (*SPSN-1: VT058873* ∩ *VT003280/FD6* and *SPSN-2: VT58873* ∩ *VT033490*) lines, which upon neuronal inhibition reduced receptivity and induced egg laying (Feng et al. 2014), for their capacity to induce the SP response upon expression of mSP. Both lines reduced receptivity and induced egg laying upon expression of mSP (Figures S4A and S4B).

Expression analysis of these two lines revealed that in addition to expression in genital tract sensory neurons (Figures S4C, S4D, S4G and S4H), they also showed expression in the brain and ventral nerve cord (Figures S4K, S4L, S4O and S4P). Intriguingly, the brain neurons labeled in SPSN-1 resembled the neurons identified by *SPR8* ∩ *FD6* (Figure 4G).

To determine whether SPSN neurons could overlap with expression of *SPR*, *dsx* and *fru* we analysed co-expression with *ocelliless* (*VT05573*), *Gyc76C* (*VT033490*) and *CG31637* (*FD6*), which are the genes where the enhancers of the *split-GAL4* lines originate, in the single cell brain atlas (Li et al. 2022). *CG31637* co-expressed in many cells with *SPR* and *fru*, but only few cells with *dsx* (Figures S6A-S6C). Expression of *ocelliless* with *SPR* and *fru* is broad, while only one neuron expressed with and *Gyc76C* in the brain (Figures S6D, S6E, S6G and S6H). Expression of *ocelliless* with *dsx* is restricted to two neurons, and no overlap was detected with *Gyc76C* in the brain Figures S6F-S6I).

When we analysed *split-GAL4* combinations of SPSN (VT058873, the common line in the SPSN1 and 2 lines) with *SPR8*, *fru11/12* and *dsx*, we observed full response to *mSP* expression for the intersection with *SPR8* and *fru11/12,* and a partial response for the *SPSN* ∩ *dsx* intersection (Figure S7A and S7B). Intriguingly, all of these *split-Gal4* combinations expressed in few neurons in the brain, the VNC and genital tract neurons except for *VY058873* ∩ *fru11/12* (Figure S7C-S7N).

We then restricted expression of mSP and induction of neuronal activity to the head with these splitGAL4 combinations using *FRTstop* casettes and *otdflp*. In this set up, we can induce PMRs from mSP expression or neuronal activation from TrpA1 expression with the *VY058873* ∩ *SPR8* and *dsx* combination, but not with the *fru11/12* combination (Suppl Fig S7O and S7P). For the VT058873 ∩fru11/12 intersection PMR inducing neurons likely reside in the VNC.

### *ppk* neurons do not intersect with SPR, fru, dsx and FD6 neurons in inducing PMRs by mSP

Expression of *UASmSP* using a *GAL4* driven by a promoter fragment of the *ppk* gene can also induce PMRs (Figures S8A and S8B) (Hasemeyer et al. 2009; Yang et al. 2009). The complement of neurons labeled with *ppkGAL4* consists of at least two populations including prominently sensory neurons, but also eight interneurons in the central brain (Nallasivan et al. 2021). These brain neurons show severe developmental defects in SP-insensitive *Nup54* mutant alleles, but they receive inhibitory input from sensory neurons (Nallasivan et al. 2021).

To evaluate whether *ppkGAL4* neurons are part of the previously identified expression patterns, we intersected them by crossing *GAL4-AD* lines *SPR8*, *SPR12* and *fru11/12* and the pan-neural *nSybAD* with a *ppk GAL4-DBD* line containing the previously used 3 kb promoter fragment (Grueber et al. 2003; Seidner et al. 2015; Riabinina et al. 2019). Surprisingly, none of these *split-GAL4* combinations reduced female receptivity or increased egg laying (Figures S8A and S8B, and Figures S8A and S8B).

Few GFP expressing neurons were detected in the brain for the *nSyb* ∩ *ppk* and the *fru11/12* ∩ *ppk* intersection (Figures S8C-S8F) or abdominal ganglion (Figures S8G-S8J). For the *nSyb* ∩ *ppk* and the *SPR8* ∩ *ppk* intersection we detected GFP expression in genital tract sensory neurons (Figures S8O and S8P), but not for the other combinations (Figures S8K and S8R).

Inhibiting or activating neurons with these split-Gal4 combinations did not reduce receptivity or induce egg laying (Figures S8L-S8O). How exactly *ppk* neurons labeled with *ppkGAL4* impact on PMRs, however, needs to be further evaluated in follow-up studies. Moreover, if genetical tract neurons were SP target sites, an SP response would have been expected for the *nSyb* ∩ *ppk* intersection, which we did not observe.

### Female post-mating neuronal circuitry contains neurons that reduce receptivity without inducing oviposition in response to mSP

A number of additional *split-GAL4* combinations with restricted expression have been identified that play a role in female reproductive behaviors (Wang et al. 2020a; Wang et al. 2020b; Wang et al. 2021). These lines express in a subset of *dsx* expressing neurons (*pC1-SS1*), in oviposition descending neurons (*oviDN-SS1* and 2), in oviposition excitatory neurons (*oviEN-SS1* and 2), in oviposition inhibitory neurons (*oviIN-SS1* and 2), and in vaginal plate opening neurons (*vpoDN-SS1*, also termed ovipositor extrusion/rejection behavior neurons (Aigaki et al. 1991; Soller et al. 2006)). When we analyzed these lines for a response to mSP expression, receptivity was reduced from mSP expression in *oviEN-SS2, oviN-SS1* and *vpoDN-SS1* neurons, but no egg laying was induced from mSP expression in any of these neurons (Figure S9A and S9B).

In genital tract neurons, *OviDN-SS1s, OviEN-SS1, OviIN-SS1* and *vpoDNs* express, but *OviDN-SS1s* and *OviEN-SS1* express weakly (Figure S9C and S9J).

### Interference of neuronal activity in SPRINz reveals regulatory hierarchy

Both inhibitory and activating neurons have been attributed to impact on PMRs (Kvitsiani and Dickson 2006; Yapici et al. 2008; Rezaval et al. 2012). These neurons seem to be part of intersecting circuitry as general inhibition of *ppkGAL4* neurons by tetanus toxin (TNT) only partially blocks the SP response in contrast to inhibition of *ppkGAL4* neurons in the brain alone (Nallasivan et al. 2021).

When we inhibited neuronal activity by expression of TNT (Sweeney et al. 1995), we observed a significant reduction of receptivity for all *split-Gal4* combinations, though only partially for inhibition in *fru11/12* ∩ *FD6* neurons. Likewise, all *split-Gal4* combinations induced a significant increase in egg laying (Figures S10A and S10B). Ablation of these neurons by expression of apoptosis inducing *reaper* and *hid* genes essentially replicated the results from neuronal inhibition indicating that SPR target neurons are modulatory and are not part of motor circuits because females laid eggs and performed normally in receptivity assays (Figures S10C and S10D).

To evaluate the composition of the intersected expression patters into inhibitory and activating neurons we also expressed the *Bacillus halodurans* sodium channel (NaChBac) (Feng et al. 2014) to activate all of the intersected neurons. Here, we found a significant reduction of receptivity for four of the five *split-GAL4* combinations, though only partially for activation of *SPR8* ∩ *dsx* neurons (Figure S10E). Activating *fru11/12* ∩ *FD6* neurons did not reduce receptivity (Figure S10E). Likewise, we found the same pattern for the induction of egg laying (Figure S10F). Four of the five *split-GAL4* combinations induced a significant increase which was only partial in *SPR8* ∩ *dsx* neurons and no egg laying was induced by activating *fru11/12* ∩ *FD6* neurons.

Essentially, these results are consistent with previous findings that inhibitory neurons prevail (Nallasivan et al. 2021), possibly as input from trunk neurons as found for *ppk* expressing neurons.

### mSP responsive neurons operate in higher order sensory processing in the brain

With the split-GAL4 approach we identified five distinct neuronal sub-types that can induce PMRs. To find out whether these neurons receive input from distinct entry points in the brain and to identify the target neurons of these mSP responsive neurons, we used the *retro-* and *trans*-Tango technique to specifically activate reporter gene expression in up- and down-stream neurons (Talay et al. 2017; Sorkaç et al. 2023)(Figures 7A-O).

**Figure 7:**
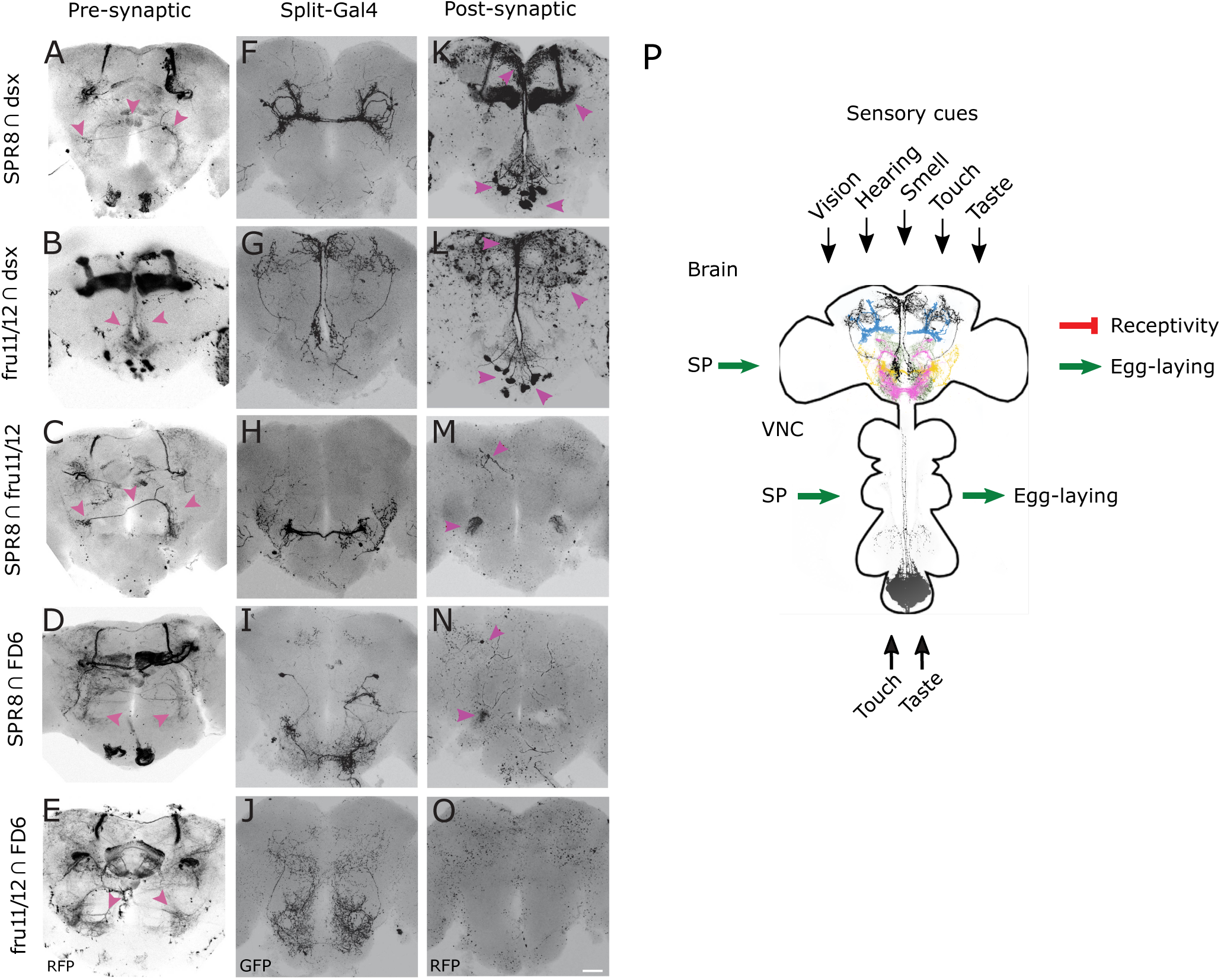
*retro-* and *trans*-Tango identification of pre- and post-synaptic neurons of SP target neurons reveals higher order neuronal input canalized into shared output circuitries. **A-O)** Representative adult female brains expressing *QUAST tomato3xHA retro-*Tango (left, A-E), *UAS myrGFP* (middle, F-J) and *QUAST tomato3xHA trans-*Tango (right, K-O) in *SPR8* ∩ *dsx, fru11/12* ∩ *dsx, SPR8* ∩ *fru11/12, SPR8* ∩ *FD6* and *fru11/12* ∩ *FD6* split-*GAL4s*. The presynaptic (A-E, left), *split-GAL4* (F-J, middle) and postsynaptic (K-O, right) neuronal circuitries are shown in an inverted grey background. Arrows (magenta) indicate neurons and their corresponding projections in different regions in female brain. The scale bar shown in O is 50 μm. **P)** Model for the SP induced post-mating response. SP interferes with interpretation of sensory cues, e.g. vison, hearing, smell, taste, and touch at distinct sites in the brain indicated by higher order projections revealed by intersectional expression in the following patterns: *SPR8* ∩ *dsx* (blue)*, fru11/12* ∩ *dsx* (black)*, SPR8* ∩ *fru11/12* (yellow)*, SPR8* ∩ *FD6* (pink) and *fru11/12* ∩ *FD6* (olive). and VNC (*fru11/12* ∩ *dsx*) during higher order neuronal processing.

In the brain, the *retro*-Tango analysis did not identify primary sensory neurons, but higher order neurons in the central brain in all five *split-GAL4* combinations (Fig 7A-E). In addition, neurons in the suboesophagal ganglion were marked from *SPR8* intersections with *dsx* and *FD6*, and in *dsx* ∩ *fru11/12*. In *dsx* ∩ *fru11/12*, neurons in the optic lobe (medulla) were marked. In addition, a strong signal was observed in all five *split-GAL4* combinations in the mushroom bodies (Figs 7A-E). Although mushroom bodies are dispensable for PMRs (Fleischmann et al. 2001) their connection to SP target neurons indicates an experience dependent component of PMRs.

The *trans*-Tango analysis identified a subset of neurons with cell bodies in the suboesophageal ganglion with projections to the *pars intercerebralis* for *SPR8* ∩ *dsx* and *fru11/12* ∩ *dsx* neurons (Figures 7K and 7L). For *SPR8* ∩ *fru11/12* and *SPR8* ∩ *FD6* neurons common target neurons were found in the antennal mechanosensory and motor centre (AMMC) region with a single neuron identified near the mushroom body region (Figures 7M and 7N) (Ishimoto and Kamikouchi 2021). For *fru11/12* ∩ *FD6* no obvious targets were identified in the central brain (Figure 7O).

In the VNC, the *trans*-Tango analysis showed post-synaptic targets within the abdominal ganglion with all five *split-GAL4* combinations indicating an interconnected neuronal network (Figure S11A-S11O), which needs to be elaborated in detail. In the genital tract, no post-synaptic targets were detected indicating that these are afferent neurons integrating sensory input (Figure S11P-S11AD).

Taken together, circuitries identified via *retro-* and *trans-*Tango place SP target neurons at the interface of sensory processing interneurons connecting to two commonly shared post-synaptic processing neuronal populations in the brain. Hence, our data indicate that SP interferes with sensory input processing from multiple modalities that are canalized to higher order processing centres to generate a behavioural output.

## Discussion

Much has been learned about the neuronal circuitry governing reproductive behaviors in *Drosophila* from interfering with neuronal activity in few neurons selected by intersectional expression using *split-GAL4* (Wang et al. 2020a; Wang et al. 2020b; Wang et al. 2021). However, how sex-peptide signaling as main inducer of the post-mating response, prominently consisting of refractoriness to re-mate and induction of egg laying, is integrated in this circuitry is not completely understood (Haussmann et al. 2013).

Here, we addressed this gap by identifying regulatory regions in *SPR*, *fru* and *dsx* genes driving membrane-tethered expression of SP in subsets of neurons to delineate SP targets to very few neurons in the central brain and the ventral nerve cord by intersectional expression. Consistent with previous analysis describing multiple pathways for the SP response (Haussmann et al. 2013), we find five distinct populations of interneurons in the central brain directing PMRs. In SP target neurons in the central brain, SPR is essential to induce PMRs when receiving SP from males through mating. From mapping post-synaptic targets by *trans*-Tango, we identified two populations of interneurons. The architecture of this circuitry is reminiscent for processing of sensory input transmitted to central brain pattern generators for behavioral output. Hence, SP interferes at several levels for coordinating PMRs, but also leaves the female the opportunity to interfere under unfavorable conditions with specific elements of PMRs, e.g., if there is no egg laying substrate, females will still not remate (Haussmann et al. 2013). Likewise, mated females will not lay eggs despite suitable egg laying substrates if parasitoid wasps are present (Kacsoh et al. 2015). Thus, the architecture of female PMRs contrasts with male-courtship behavior consisting of a sequel of behavioral elements that once initiated will always follow stereotypically to the end culminating in mating, or start from the beginning when interrupted (Hall 1994; Greenspan and Ferveur 2000).

### SP induces PMRs via entering the hemolymph to target neurons in the central brain and ventral nerve cord

Early characterization of the SP signaling cascade demonstrated induction of PMRs from various other sources than mating including transgenic secretion from the fat body, expression as membrane-tethered form on neurons or injection of synthetic peptide into the hemolymph (Chen et al. 1988; Aigaki et al. 1991; Schmidt et al. 1993; Nakayama et al. 1997). Likewise, SP is detected in the hemolymph after mating at a PMR inducing concentration (Haussmann et al. 2013). Moreover, PMRs are induced faster, when SP is injected compared to induction by mating (Haussmann et al. 2013). This delay, however, is not attributed to sperm binding of SP as it is unchanged after mating with spermless males. These results suggest that SP reaches its targets through entering the circulatory system to target neurons and contrasts a previously proposed model favoring genital tract neurons as SP sensors from the lumen of the genital tract (Hasemeyer et al. 2009; Yang et al. 2009; Rezaval et al. 2012).

In further support of the internalization model, we identified *GAL4* drivers that express mSP in genital tract neurons, but do not induce PMRs. Also, *SPR12* does not express in genital tract neurons, but induces egg laying by expression of mSP. Moreover, expression of mSP predominantly in the trunk (including all genital tract sensory neurons), only induces egg laying, but does not change receptivity. Likewise, expression of mSP specifically in the brain (*SPR8* ∩ *dsx*) can reduce receptivity and induce egg laying indistinguishable from mated females.

A *ppkGAL4* line generated by P-element mediated transformation can induce PMRs by expression of *UAS mSP* (Grueber et al. 2003). The same promoter fragment fused to a *GAL4 DBD* and inserted by phiC31 integration into a landing site intersected with pan-neural *nSyb AD* line (Seidner et al. 2015; Riabinina et al. 2019), however, does not induce a SP response despite being expressed in genital tract neurons. We found that the *ppkGAL4* expresses in a few neurons in the brain and VNC (Nallasivan et al. 2021), but this expression is absent in *nSyb* ∩ *ppk* intersection. Likely, the *ppkGAL4* construct is inserted in a locus that contains an enhancer that drives expression in SP target neurons.

These results are in strong favor for SP entering the hemolymph to target neurons in the ventral nerve cord for inducing egg laying, and in the central brain for reducing receptivity and inducing egg laying (Haussmann et al. 2013).

### Integration of SP signaling into the circuitry directing reproductive behaviours

Reduction of receptivity and induction of egg laying are both induced by the same critical concentration of injected SP (Schmidt et al. 1993; Haussmann et al. 2013) initially suggesting a simple on/off system for PMRs likely initiated from a small population of neurons. However, such model would not allow to split the SP response into individual PMR components by expression of mSP.

Here, we identified several *GAL4* drivers, that can induce only egg laying (*SPR12*, *FD3*, *FD4* and *tsh GAL4*), but do not reduce receptivity, and others that can only reduce receptivity (*oviEN-SS2, oviIN-SS1* and *vpoDN-SS1*), but do not induce egg laying. Strikingly *tshGAL4*, that expresses predominantly in the trunk only affects egg laying suggesting a role for the abdominal ganglion in egg laying. Moreover, all of the *split-GAL4* combinations affecting egg laying express in the abdominal ganglion and *dsx* neurons in the abdominal ganglion have been identified to induce egg laying (Rezaval et al. 2012; Zhou et al. 2014). Hence, this neuronal structure has a key role in regulating egg laying. Since more than a single neuronal population seems to direct egg laying, further high-resolution mapping is required to identify individual neuronal population within the abdominal ganglion (Jang et al. 2017; Oliveira-Ferreira et al. 2023).

Since *tshGAL4* only induces egg laying, neurons in the brain must direct reduction of receptivity. Through intersectional expression in combination with head-specific expression of *otdflp*, we could express mSP only in the brain by FLP mediated brain-specific excision of a stop cassette. We observed a significant reduction in receptivity for all five intersections tested, but for four the response is only partial likely due to the inefficiency of FLP mediated recombination.

Moreover, brain neurons can also induce egg laying when *SPR8* is intersected with *dsx*, and to some extent also from *SPR8* intersection with *fru11/12*. Due to the inefficiency of FLP mediated recombination, however, this is likely an underestimate and solving this issue requires development of more robust tools.

In any case, however, our results show that PMRs can be induced from mSP expression from several sites suggesting interference with processing of sensory information at the level of interneurons. In particular, *SPR8* ∩ *fru11/1*2 neurons resemble auditory AMMC-B2 neurons involved in processing of information of the male love song (Yamada et al. 2018). Likewise, *SPR8* ∩ *dsx* neurons seem to overlap with dimorphic *dsx* pCL2 interneurons that are part of the 26 neurons constituting the pC2 neuronal population involved in courtship song sensing, mating acceptance and ovipositor extrusion for rejection of courting males (Kimura et al. 2015; Deutsch et al. 2019; Wang et al. 2020a). The *SPR8* ∩ *FD6* neurons resemble dopaminergic *fru* P1 neurons involved in courtship and the *fru11/12* ∩ *dsx* neurons seem to overlap with *dsx* pCd and neuropeptide F neurons involved in courtship (Zhang et al. 2021). In females, pC1d neurons have been linked to aggression (Deutsch et al. 2020; Schretter et al. 2020). The *fru11/12* ∩ *FD6* neurons resemble a class of gustatory pheromone sensing neurons (Sakurai et al. 2013). Although we likely have not identified all SP sensing neurons, our resources will provide a handle to future exploration of the details of this neuronal circuitry incorporating SP signaling for inducing PMRs.

## Conclusions

We have identified distinct SP sensing neurons in the central brain and the ventral nerve cord. Since these five different SP sensing neuronal populations in the central brain converge into two target sites, our data suggest a model (Figure 7P), whereby SP signaling interferes with integration of sensory input. Independent interference with different sensory modalities opts for the female to counteract male manipulation at the level of perception of individual sensory cues to adapt to varying physiological and environmental conditions to maximize reproductive success.

## Acknowledgments

We thank T. Aigaki, G. Barnea, P. Soba, W.J. Joiner, B. Dickson, S. Goodwin, C. Rezaval, D. Anderson, J.J. Hodge, A. Hidalgo, S. Collier, O. Raibinina, the Bloomington stock center, the Vienna Drosophila RNAi Center for flies, T. Aigaki and W.J. Joiner for plasmids, the University of Cambridge Department of Genetics Fly Facility and FlyORF for injections, D. Scocchia for help with PCR and, I.U. Haussmann, Y.J. Kim, J.C. Billeter and J-R Martin for comments on the manuscript. We acknowledge funding by the Biotechnology and Biological Science Research Council to MS.

## Author contributions

MS conceived and directed the project. MPN performed genetic experiments and imaging. MS and SS performed genetic and DS imaging experiments. MPN and MS analyzed data. MS wrote the manuscript with support from MPN. All authors read and approved the final manuscript.

## Data availability

Brain and VNC images for splitGal4 combinations of SP Response Inducing Neurons have been deposited in Virtual Fly Brain and will be published under the following accession numbers: VFB_x0000000-9. All data generated or analysed during this study are included in the paper and supplementary files; source data files are provided for all figures.

## Declarations

### Ethics approval and consent to participate

Not applicable.

### Consent for publication

Not applicable.

## Materials and methods

### Key resources table

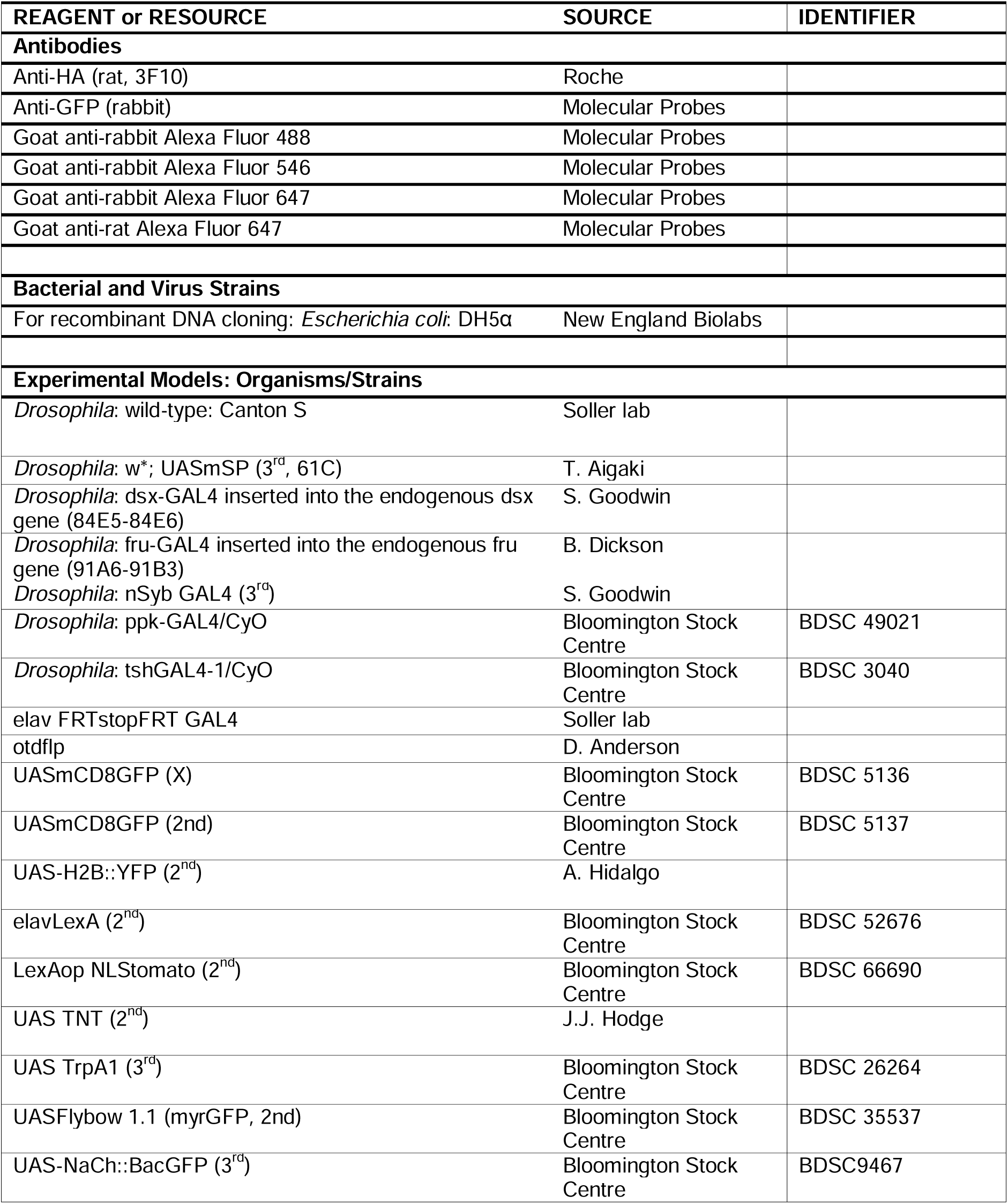
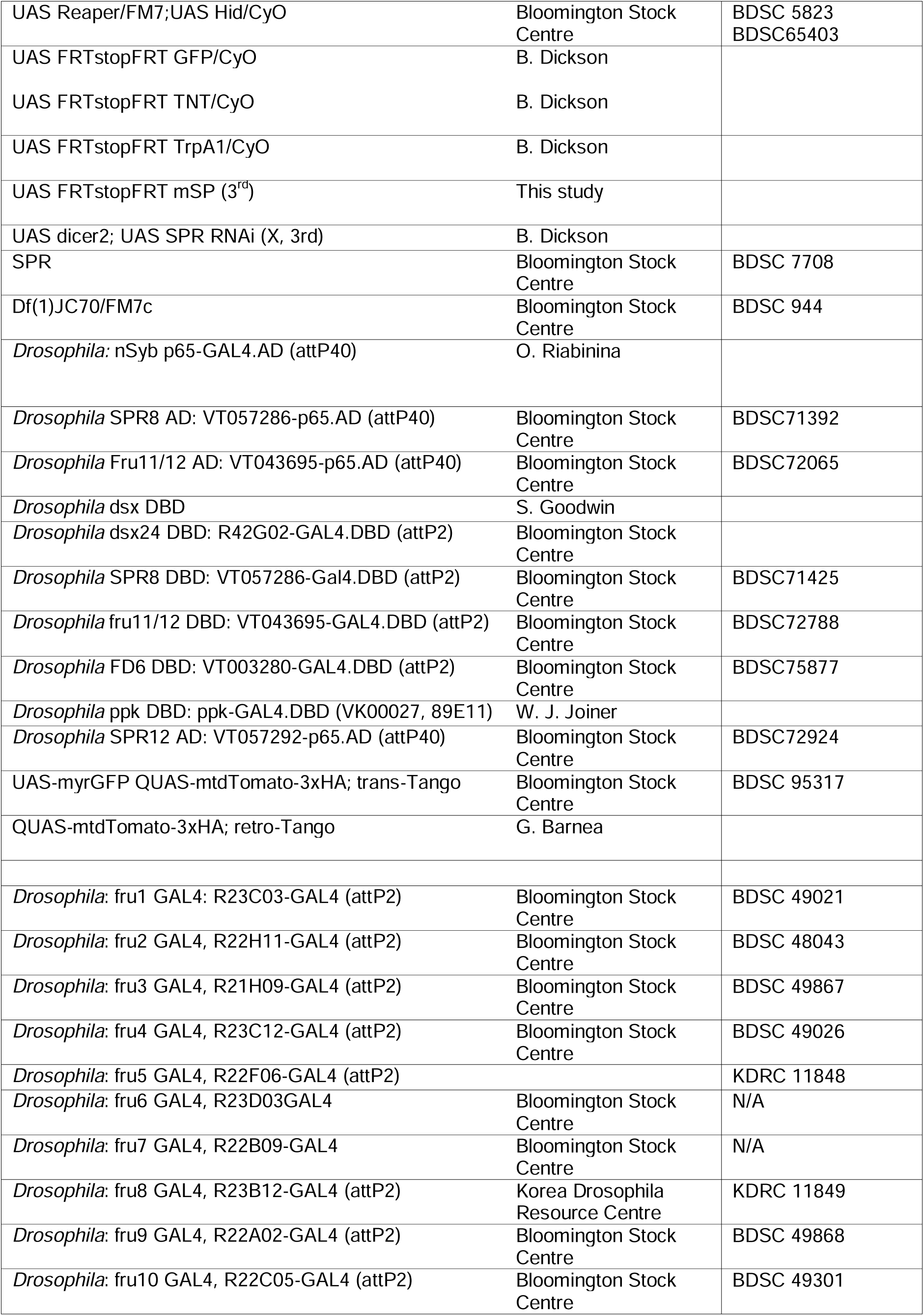
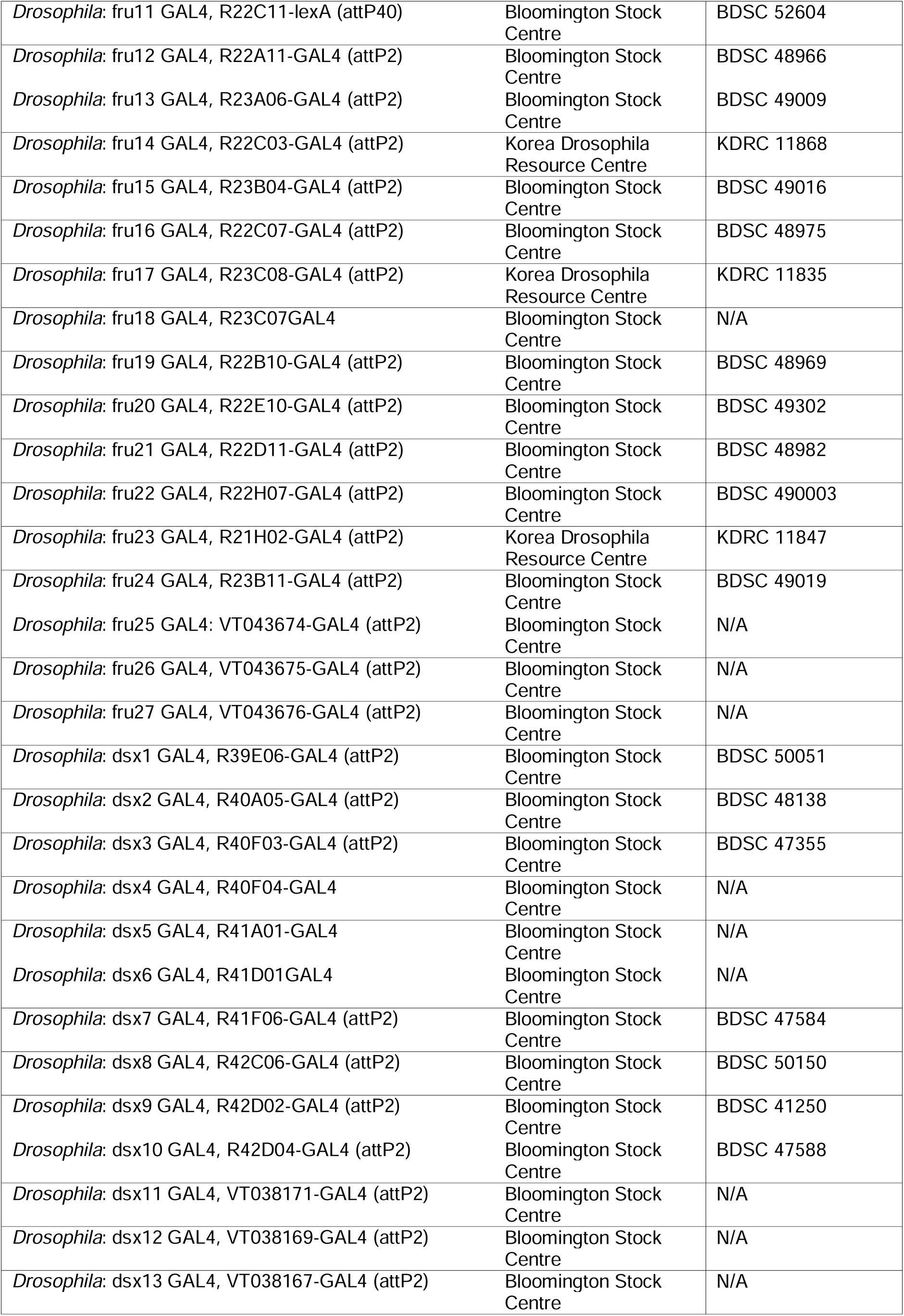
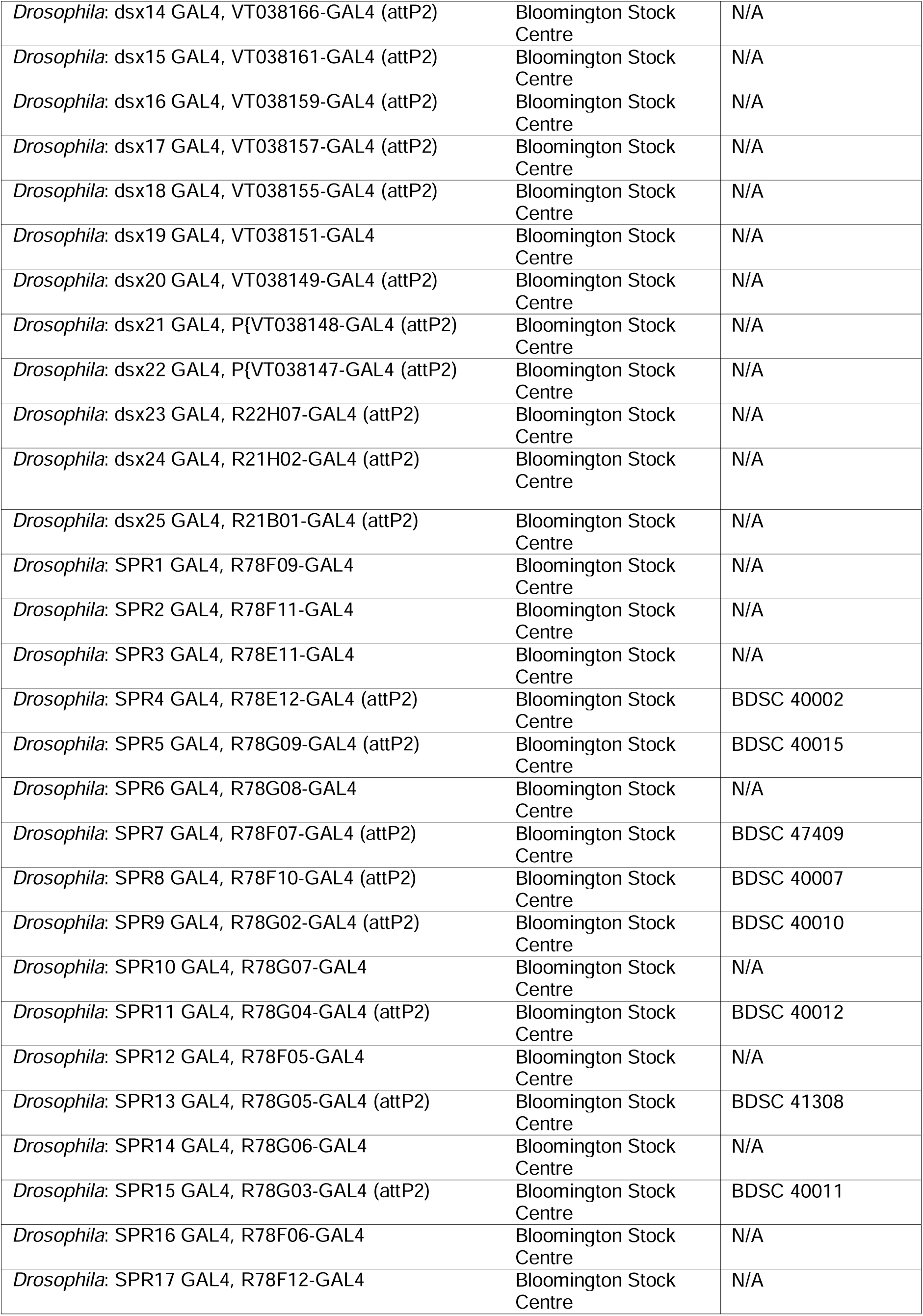
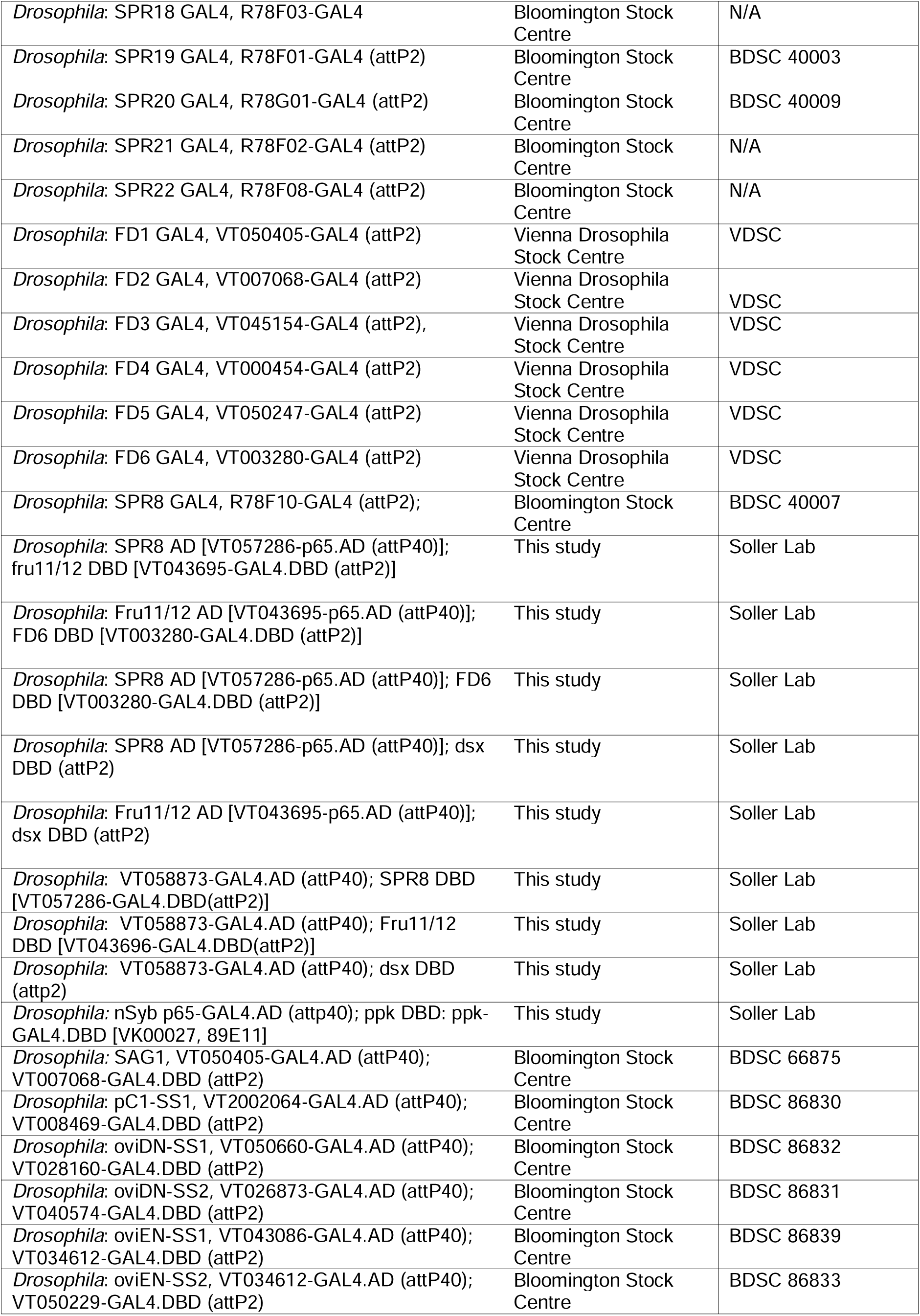
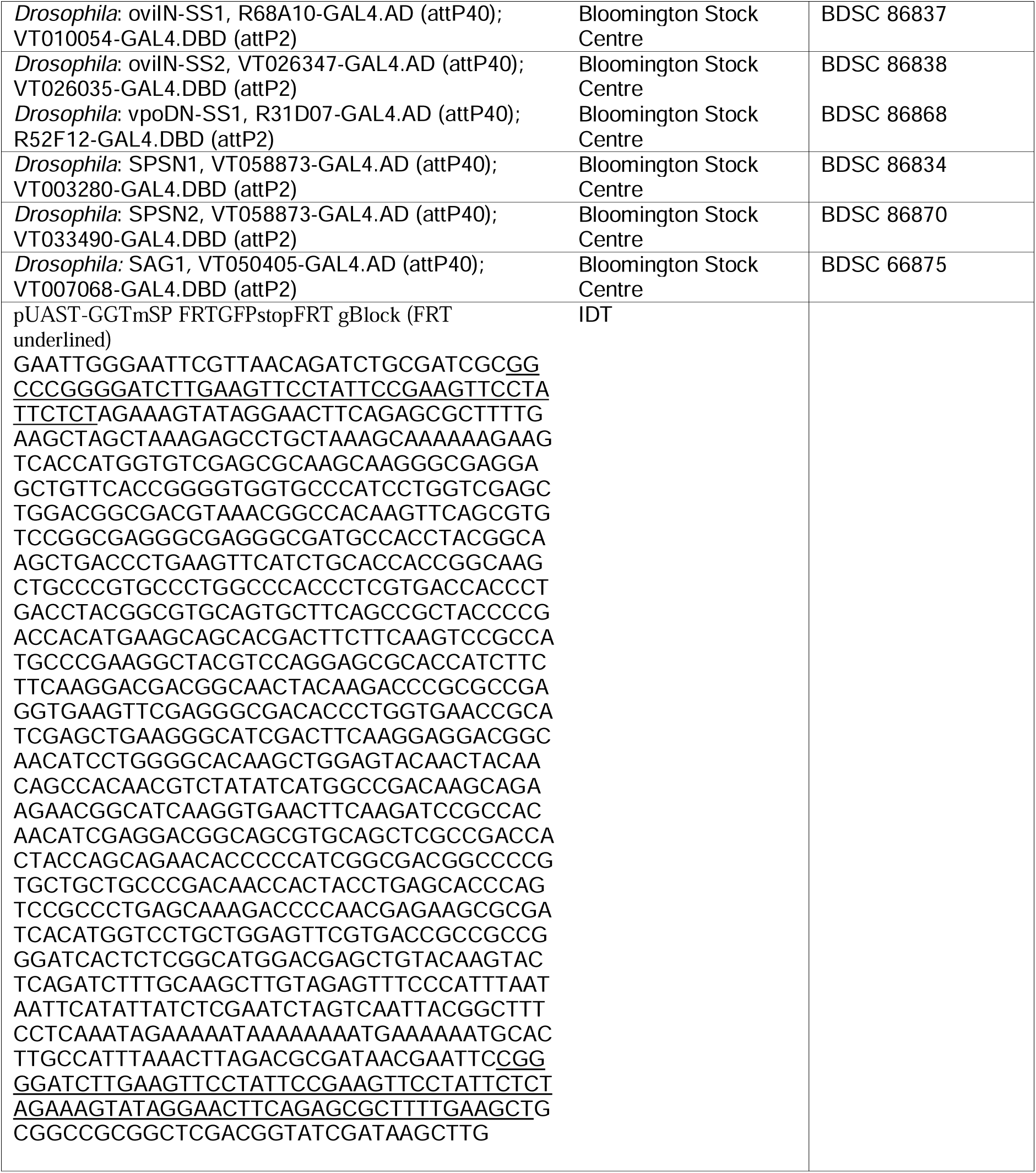

### Fly strains and husbandry

Flies were kept on standard cornmeal-agar food (1%industrial-grade agar, 2.1% dried yeast, 8.6% dextrose, 9.7% cornmeal and 0.25% Nipagin, all in (w/v)) in a 12 h light : 12 h dark cycle. Propionic acid was omitted from fly food as acidity affects egg laying (Gou et al. 2014). Genetic crosses were done in vials and kept at low density to ensure larvae were not competing for food and if necessary, additional live yeast was added. For all behavioral assays, virgin and mated Canton-S were used as controls. Virgin females, e.g. from crosses of *GAL4* with *UASmSP*, were collected after emergence within a 5 h window and well-fed with live yeast sprinkled on food for maximum egg production and allowed to sexually mature (3– 5 days).

To recombine 2^nd^ chromosome inserts for *splitGAL4AD* (*attP40*) and 3^rd^ chromosome *splitGAL4DBD* (*attP2*), standard genetic crossing schemes were used and final stocks were balanced with CyO and TM3 Sb (combined from from ST and CT stock, see key resource list). *SplitGal4AD* and *DBD* combination lines were then crossed to *UASmSP*. For meiotic recombination, final stocks were validated by behavioral analysis for *UAS mSP*, for *flp* with *eFeG UASCD8GFP* to monitor GFP expression and for *otdflp UASstopTrpA* and *otdflp UASstopTNT* by crossing to *elavGAL4* and monitored by lethality.

For enhanced recombination with *flp*, virgin females were transferred to 30° C after eclosion and kept for 5 d at this temperature before performing the behavioral assays. For induction of neuronal activity by temperature sensitive TrpA1, females were kept at 30° C.

To make *UAS FRTstopFRT mSP*, a gBlock (IDT) stop cassette with the FRT sequences used in the eFeG plasmid (Haussmann et al. 2008) was inserted into NotI cut *pUAST-GGTmSP* (gift from T. Aigaki) by Gibson assembly. In the stop-cassette, the *FRT* sequence is followed by a *GFP* with a 3’UTR from *ewg* containing polyA site 1 from intron 6 (Haussmann et al. 2011). Flies were transformed by *P*-element mediated transgenesis and a inserts on each chromosome were established that show a robust post-mating response with *dsxflp* indistinguishable from mated females.

### Behavioral analysis

Females were examined for the main post-mating behaviors receptivity and oviposition as described previously and as follows (Soller et al. 1999; Soller et al. 2006). To generate mated females, one female and three males were added to fly vials and observed until mating and males were removed after mating. For receptivity tests, mature virgin or mated females were added to fly vials (95 mm length and 24 mm diameter) containing Canton S males with an aspirator and observed for 1 h, generally 3 females and 7 males. For these experiments, males were separated from females at least one day before the experiment. Receptivity tests were done in the afternoon with virgins, or 5-24 h after mating for controls. For oviposition, females were placed individually in fly vials in the afternoon and the number of eggs laid was counted the next day.

### Statistical analysis

Sample size was based on previous studies, non-blinded and not pre-determined by statistical methods (Soller et al. 1997; Haussmann et al. 2013; Nallasivan et al. 2021). Behavioral data are representatives of at least three replicates that were performed on three different days. Statistical analysis of behavioral experiments were performed using GraphPadPrism 9 (GraphPad by Dotmatics) using one way Anova followed by pairwise comparisons with Tukey’s test.

### Immunohistochemistry and imaging

For the analysis of adult neuronal projection from *UAS CD8GFP, UAS H2BYFP,UASmyrGFP, lexAopNLStomato* o*r QUAS mtdtomato3xHA* expressing brains, ventral nerve cords or genital tracts, tissues were dissected in PBS (137 mM NaCl, 10 mM phosphate, 2.7 mM KCl, pH7.4), fixed in 4% (w/v in PBS) paraformaldehyde for 15 minutes, washed three times in PBST (PBS with 1% BSA and 0.3% Triton-X100), then once in PBS for 10 mins, mounted in Vectashield (Vector Labs) and visualized with confocal microscopy using a Leica TCS SP8. If signals were weak, antibody *in-situ* stainings were done as described previously (Haussmann et al. 2008) for validation using rat anti-HA (MAb 3F10, 1:20; Roche), rabbit anti-GFP (Molecular Probes, 1:100) and visualized with Alexa Fluor 488 (1:250; Molecular Probes or Invitrogen), Alexa Fluor 546 (1:250; Molecular Probes or Invitrogen) or Alexa Fluor 647 (1:250; Molecular Probes or Invitrogen). For imaging, tissues were mounted in Vectashield (Vector Labs).

### Confocal microscopy and image processing

Adult tissues were scanned using a Leica SP8 confocal microscope equipped with a set of fluorescent filters and hybrid detector (HyD). Adult brains were scanned using a 40x HC PL APO 40x/1.30 lens with oil, 1024 x 1024 resolution and 0.96 µm Z-step. VNC and genital tracts were scanned using a HC PL APO CS2 20X/0.75 with oil, 1024 x 1024 resolution and 0.96 µm Z-step. Images were obtained using Leica Application Suite X (LAS X) imaging acquisition software. Raw data files were in LIF format and were processed using FIJI.

For high resolution mapping neurons were identified in the virtual fly brain based on registered *GAL4* expression and traces retrieved for modelling (Scheffer et al. 2020; Phelps et al. 2021; Galili et al. 2022).

## Supplementary information

**Supplementary Figure S1: Analysis of head and trunk expression lines.**

**A-F)** Expression of *UAS CD8 GFP* driven *elav FRTstopFRT GAL4* restricted with *otdflp* to the head in the brain and VNC.

**G-L)** Expression of *tshGAL4 UAS H2B YFP* with neurons labeled with tomato from *elavLexA AopNLStomato* in the brain and VNC.

**Supplementary Figure S2: Expression analysis of PMR-inducing *GAL4* in the genital tract.**

**A-E)** Representative adult female genital tracts expressing *UAS CD8GFP* under the control of *SPR8*, *SPR12*, *fru11*, *fru12* and *dsx24 GAL4*, and *LexAop NLStomato* under the control of *elavLexA*. Arrows indicate genital tract sensory neurons. The insert shows expression of GFP in the genital tract sensory neurons. Scale bars shown in A and insets are 100 µm and 20 µm, respectively.

**Supplementary Figure S3: Expression analysis of non-PMR-inducing *fru9GAL4* in the genital tract.**

Representative adult female genital tract expressing *UAS CD8GFP* under the control of *fru9 GAL4*. Arrows indicate genital tract sensory neurons. The insert shows expression of GFP in the genital tract sensory neurons. Scale bars shown in C and insets are 100 µm and 20 µm, respectively.

**Supplementary Figure 4: Expression of mSP in *SPSN*, and genital tract expression *SPR* lines does not support a major role for genital tract neurons in inducing the sex peptide response.**

**A, B)** Receptivity (A) and oviposition (B) of wild type control virgin (red) and mated (orange) females, and virgin females expressing *UAS mSP* (green) under the control of *SPSN 1* and *SPSN2*, and *and SPR3* and *SPR9 GAL4* lines shown as means with standard error from three repeats for receptivity (21 females per repeat) by counting the number of females mating within a 1 h period or for oviposition by counting the eggs laid within 18 hours from 30 females. Statistically significant differences from ANOVA post-hoc comparison are indicated by different letters (p<0.0001).

**C-J)** Representative genital tracts labelled with UAS CD8 GFP and genital tract neurons labelled with UAS H2BYFP and *elavLexA AopNLStomato*.

**K-R)** Adult female brains (K-N) and ventral nerve cords (VNC. O-R) expressing *UAS CD8GFP*. Scale bars shown in H and M are 50 µm and 100 µm, respectively.

**Supplementary Figure S5: Expression analysis of split-GAL4 in the genital tract.**

**A-E)** Representative adult female genital tracts expressing *UAS CD8GFP* under the control of *SPR8* ∩ *fru11/12, SPR8* ∩ *dsx, SPR8* ∩ *FD6, fru11/12* ∩ *dsx* and *fru11/12* ∩ *FD6 split-GAL4* intersectional patterns. The scale bar shown in E is 100 µm.

**Supplementary Figure S6: Expression analysis of split-GAL4 in the genital tract.**

**A-C)** Visualisation of single cell expression for *CG31637* intersected with *SPR*, *fru* and *dsx*. **D-F)** Visualisation of single cell expression for *ocelliless* intersected with *SPR*, *fru* and *dsx*. **G-I)** Visualisation of single cell expression for *Gyc76c* intersected with *SPR*, *fru* and *dsx*.

**Supplementary Figure 7: Expression of mSP in *SPSN VT058873 AD* intersected with *SPR8 DBD, fru11/12 DBD* and *dsx DBD* induces PMRs and *SPSN VT058873 AD* intersected with *SPR8 DBD* and *dsx DBD* sense SP in the brain.**

**A, B)** Receptivity (A) and oviposition (B) of wild type control virgin (red) and mated (orange) females, and virgin females expressing *UAS mSP* (green) under the control of *VT058873* ∩ *SPR8, VT058873* ∩ *fru11/12 and VT058873* ∩ *dsx* shown as means with standard error from three repeats for receptivity (21 females per repeat) by counting the number of females mating within a 1 h period or for oviposition by counting the eggs laid within 18 hours from 30 females. Statistically significant differences from ANOVA post-hoc comparison are indicated by different letters (p<0.0001).

**C-H)** Adult female brains (C-E) and ventral nerve cords (VNC, F-H) expressing *UAS CD8GFP*. Scale bars shown in H and M are 50 µm and 100 µm, respectively.

**I-N)** Representative genital tracts labelled with UAS CD8 GFP and genital tract neurons labelled with *UAS H2BYFP* and *elavLexA AopNLStomato*.

**O, P)** Receptivity (O) and oviposition (P) of wild type control virgin (red) and mated (orange) females, and virgin females expressing *UAS FRTGFPstopFRTmSP* (grey) and *UAS FRTGFPstopFRTTrpA1* (purple) under the control of *split-GAL4* intersecting *VT058873* ∩ *SPR8, VT058873* ∩ *fru11/12 and VT058873* ∩ *dsx* patterns with brain-specific FRT-mediated recombination by *otdflp* shown as means with standard error from three repeats for receptivity (21 females per repeat) by counting the number of females mating within a 1 h period or for oviposition by counting the eggs laid within 18 hours from 30 females. Statistically significant differences from ANOVA post-hoc comparison are indicated by different letters (p<0.0001).

**Supplementary Figure S8: *ppk* is not part of the *SPR8, SPR12 and fru11/12* PMR-inducing neuronal circuitry**

**A, B)** Receptivity (A) and oviposition (B) of wild type control virgin (red) and mated (orange) females, and virgin females expressing *UAS mSP* (green) under the control of *GAL4* in *ppk* or in *nSyb* ∩ *ppk, SPR8* ∩ *ppk, SPR12* ∩ *ppk,* and *fru11/12* ∩ *ppk* patterns shown as means with standard error from three repeats for receptivity (21 females per repeat) by counting the number of females mating within a 1 h period or for oviposition by counting the eggs laid within 18 hours from 30 females. Statistically significant differences from ANOVA post-hoc comparison are indicated by different letters (p<0.0001).

**C-R)** Representative adult female brains, ventral nerve cords (VNC) and genital tracts expressing *UAS CD8GFP* under the control of *UAS* by *nSyb* ∩ *ppk, SPR8* ∩ *ppk, SPR12* ∩ *ppk,* and *fru11/12* ∩ *ppk*. Scale bars shown in E are 50 µm and in H and K are 100 µm, respectively.

**S-V)** Receptivity (S, T) and oviposition (U, V) of wild type control virgin (red) and mated (orange) females, and virgin females expressing either *UAS TNT* (azure) or *UAS NaChBac* (brown) to inhibit or activate neurons in *SPR8* ∩ *ppk, SPR12* ∩ *ppk,* and *fru11/12* ∩ *ppk* patterns shown as means with standard error from three repeats for receptivity (21 females per repeat) by counting the number of females mating within a 1 h period or for oviposition by counting the eggs laid within 18 hours from 30 females. Statistically significant differences from ANOVA post-hoc comparison are indicated by different letters (p<0.001 for b, and p<0.01 for c in L and N).

**Supplementary Figure 9: Expression of mSP in female reproductive behavior regulating neuron *split-GAL4* lines.**

**A, B)** Receptivity (A) and oviposition (B) of wild type control virgin (red) and mated (orange) females, and virgin females expressing *UAS mSP* (green) under the control of *pC1-SS1, oviDN-SS1 and 2, oviEN-SS1 and 2, oviIN-SS1 and 2, and vpoDN-SS1* shown as means with standard error from three repeats for receptivity (21 females per repeat) by counting the number of females mating within a 1 h period or for oviposition by counting the eggs laid within 18 hours from 30 females. Statistically significant differences from ANOVA post-hoc comparison are indicated by different letters (p<0.0001).

**C-J)** Representative genital tract neurons labelled with *UAS H2BYFP* and *elavLexA AopNLStomato*.

**Supplementary Figure S10: PMRs after neuronal inhibition, ablation or activation of distinct circuits from intersection of *SPR*, *fru*, *dsx* and *FD6* patterns in the brain and VNC.**

**A-F)** Receptivity (A, C and E) and oviposition (B, D and F) of wild type control virgin (red) and mated (orange) females, and virgin females expressing either *UAS TNT* (azure, A and B) or *UAS reaper hid* to inhibit or ablate neurons (yellow, C and D), respectively, or *UAS NaChBac* (brown, E and F) to activate neurons in *SPR8* ∩ *fru11/12, SPR8* ∩ *dsx, SPR8* ∩ *FD6, fru11/12* ∩ *dsx* and *fru11/12* ∩ *FD6 split-Gal4* patterns shown as means with standard error from three repeats for receptivity (21 females per repeat) by counting the number of females mating within a 1 h period or for oviposition by counting the eggs laid within 18 hours from 30 females. Statistically significant differences from ANOVA post-hoc comparison are indicated by letters (p≤0.0095 in A and B, p<0.0001 in C and D except p=0.016 for c in D, p<0.0001 in E and p<0.0002 in F).

**Supplementary Figure S11: *trans*-Tango identifies post-synaptic proceeding neurons of SP targets in the VNC, but not the genital tract**

**A-AD)** Representative adult female ventral nerve cords (VNC, A-O) and genital tracts (P-AD) expressing *UAS myrGFP; QUAST tomato3xHA trans-*Tango in *SPR8* ∩ *fru11/12, SPR8* ∩ *dsx, SPR8* ∩ *FD6, fru11/12* ∩ *dsx* and *fru11/12* ∩ *FD6* split-*GAL4s*. The presynaptic (A-E and P-T) and postsynaptic (F-J and U-Y) neuronal circuitries are shown in an inverted grey background and the merge is shown in colour. In the merged picture (K-O and Z-AD), the pre-synaptic and post synaptic neuronal circuitry is shown in green and magenta, respectively. Scale bars shown in O and AD are 100 μm.

