## Supplemental Figs 1-11 for "Sex-peptide targets distinct higher order processing neurons in the brain to induce the female post-mating response"

### Supp. Fig. S1

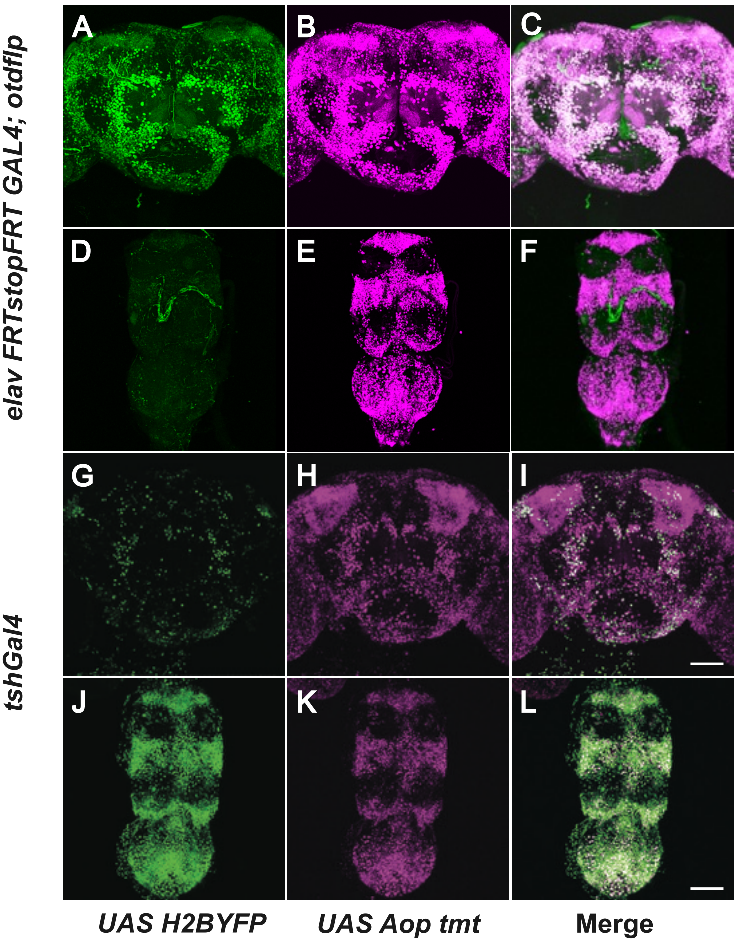
A
B
C
D
E
F
G
H
I
J
K
L
elav FRTstopFRT GAL4; otdflp
tshGal4
UAS H2BYFP
UAS Aop tmt
Merge

### Supp. Fig. S2

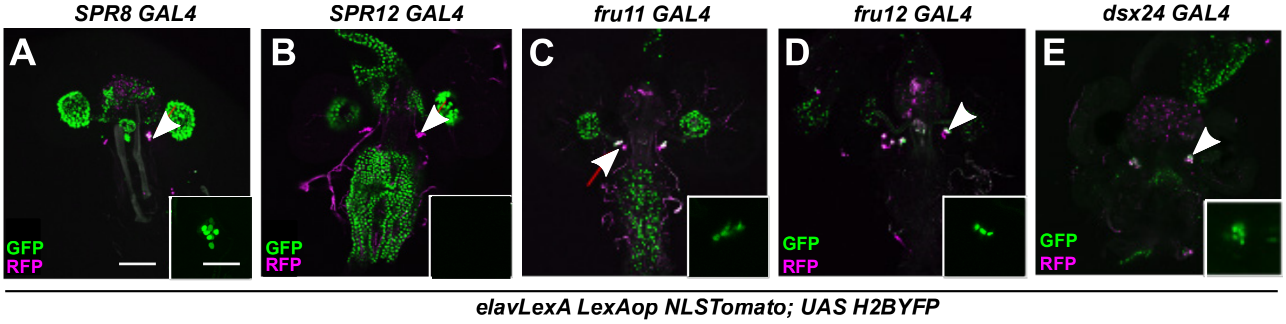
SPR8 GAL4
SPR12 GAL4
fru11 GAL4
fru12 GAL4
dsx24 GAL4
A
B
C
D
E
GFP
RFP
GFP
RFP
GFP
RFP
GFP
RFP
elavLexA LexAop NLSTomato; UAS H2BYFP

#### Supp. Fig. S3

*fru9 GAL4*

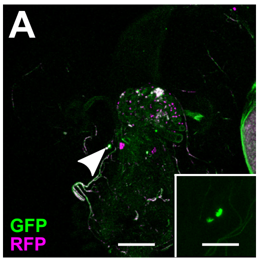


*elavLexA LexAop NLStomato ;*
*UAS H2BYFP*

### Supp. Fig. S4

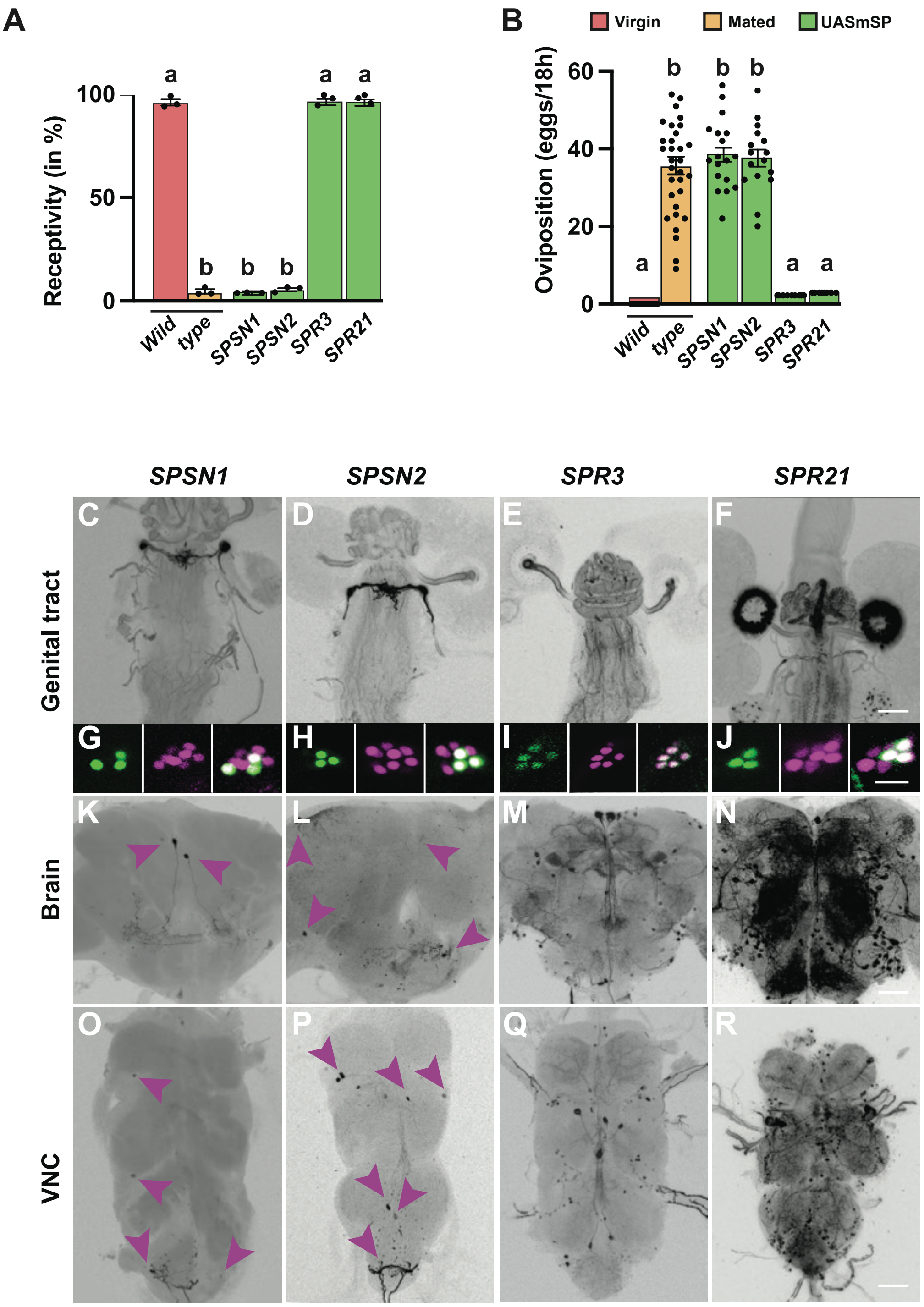
A
Receptivity (in %)
100
50
0
a
b
b
b
a
a
Wild
type
SPSN1
SPSN2
SPR3
SPR21
B
Virgin
Mated
UASmSP
Oviposition (eggs/18h)
60
40
20
0
a
b
b
b
a
a
Wild
type
SPSN1
SPSN2
SPR3
SPR21
SPSN1
SPSN2
SPR3
SPR21
Genital tract
Brain
VNC
C
D
E
F
G
H
I
J
K
L
M
N
O
P
Q
R

### Supp. Fig. S5

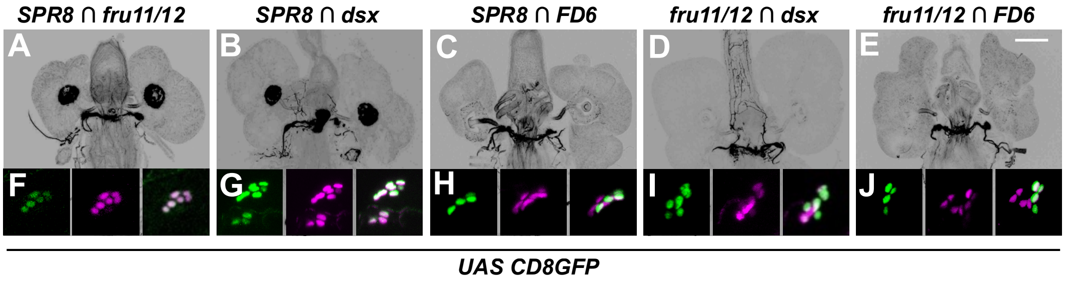

SPR8 ∩ fru11/12
SPR8 ∩ dsx
SPR8 ∩ FD6
fru11/12 ∩ dsx
fru11/12 ∩ FD6
A
B
C
D
E
F
G
H
I
J
UAS CD8GFP

### Supp. Fig. S6

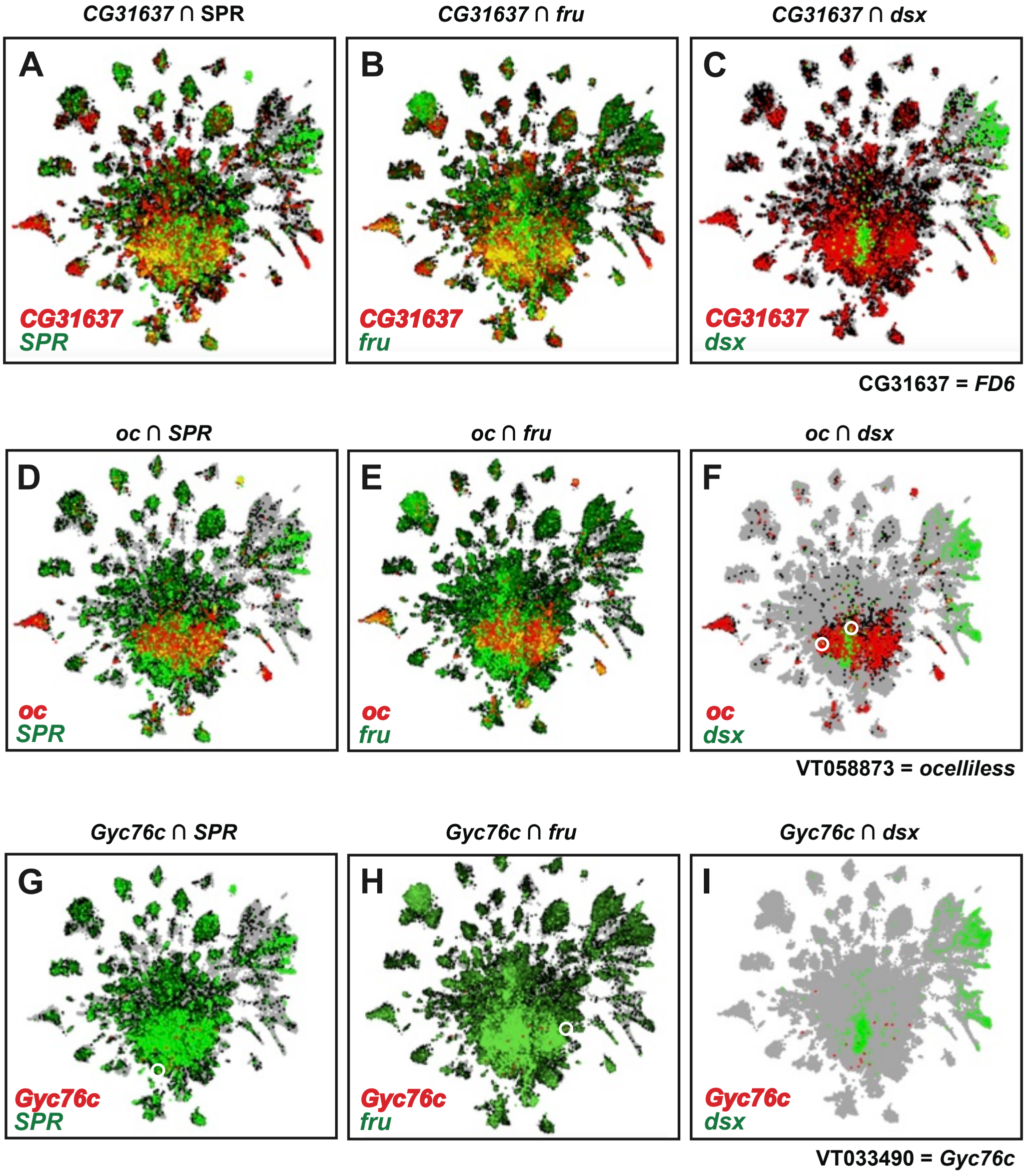

*CG31637* ∩ SPR
*CG31637* ∩ *fru*
*CG31637* ∩ *dsx*
A
B
C
CG31637
SPR
CG31637
fru
CG31637
dsx
CG31637 = *FD6*
*oc* ∩ *SPR*
*oc* ∩ *fru*
*oc* ∩ *dsx*
D
E
F
oc
SPR
oc
fru
oc
dsx
VT058873 = *ocelliless*
*Gyc76c* ∩ *SPR*
*Gyc76c* ∩ *fru*
*Gyc76c* ∩ *dsx*
G
H
I
Gyc76c
SPR
Gyc76c
fru
Gyc76c
dsx
VT033490 = *Gyc76c*

### Supp. Fig. S7

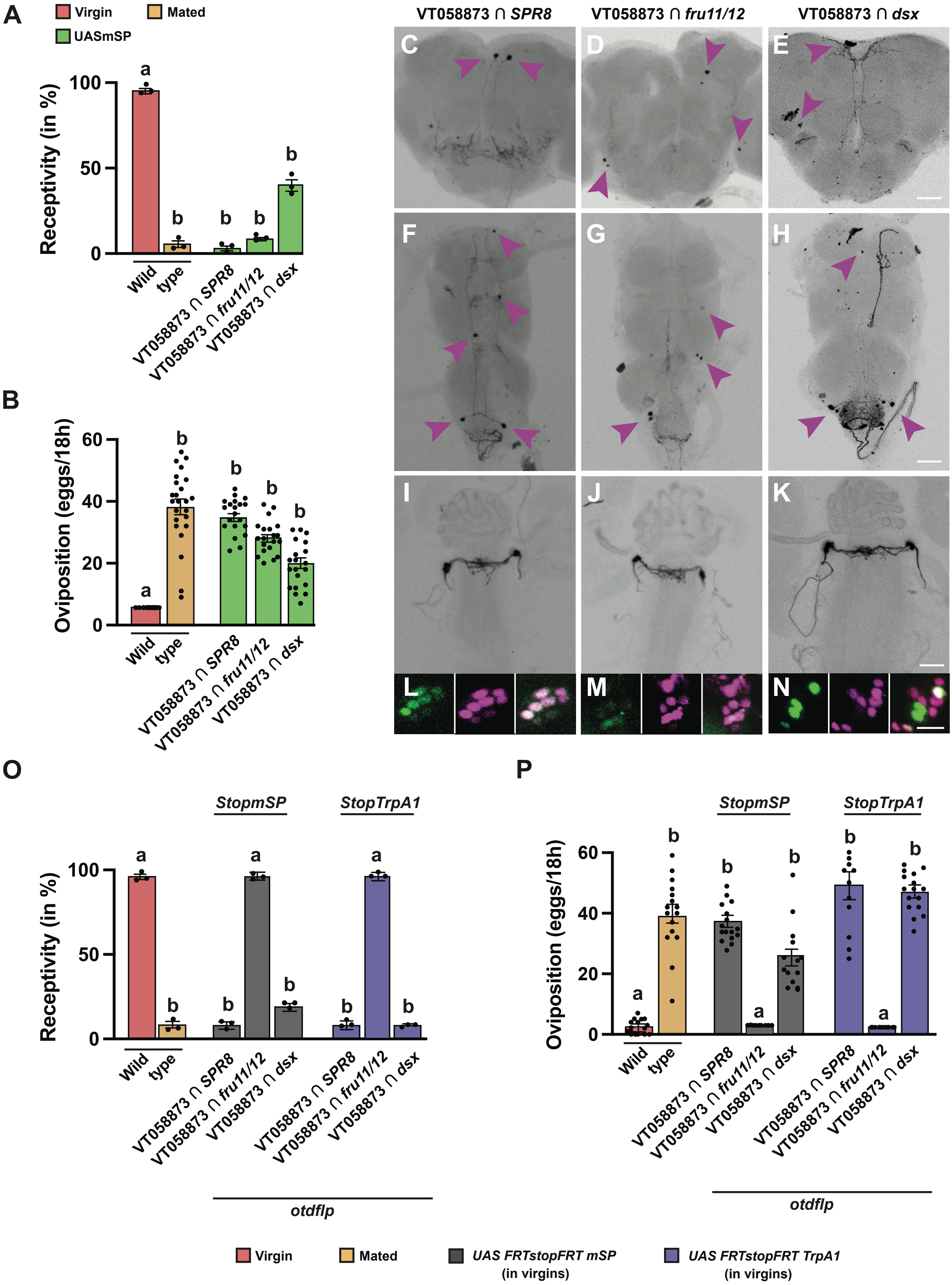

A
Virgin
Mated
UASmSP
Receptivity (in %)
Wild type
VT058873 ∩ SPR8
VT058873 ∩ fru11/12
VT058873 ∩ dsx
B
Oviposition (eggs/18h)
C D E F G H I J K L M N
O
P
StopmSP
StopTrpA1
otdflp
UAS FRTstopFRT mSP (in virgins)
UAS FRTstopFRT TrpA1 (in virgins)

### Supp. Fig. S8

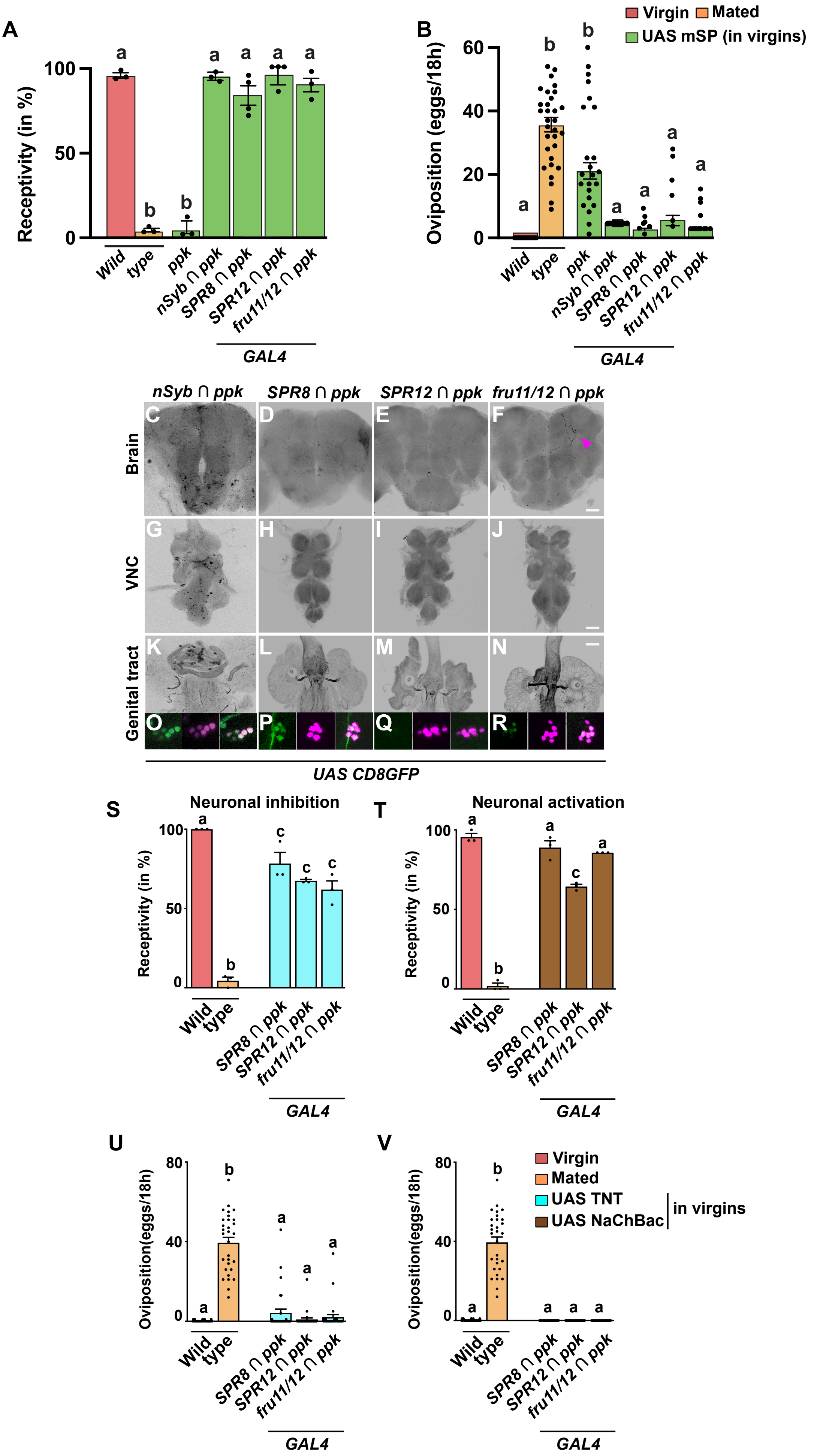

A
Receptivity (in %)
Wild type
ppk
nSyb ∩ ppk
SPR8 ∩ ppk
SPR12 ∩ ppk
fru11/12 ∩ ppk
GAL4
B
Oviposition (eggs/18h)
Virgin
Mated
UAS mSP (in virgins)
nSyb ∩ ppk
SPR8 ∩ ppk
SPR12 ∩ ppk
fru11/12 ∩ ppk
Brain
VNC
Genital tract
UAS CD8GFP
S
Neuronal inhibition
T
Neuronal activation
U
Oviposition(eggs/18h)
V
UAS TNT
UAS NaChBac
in virgins

### Supp. Fig. S9

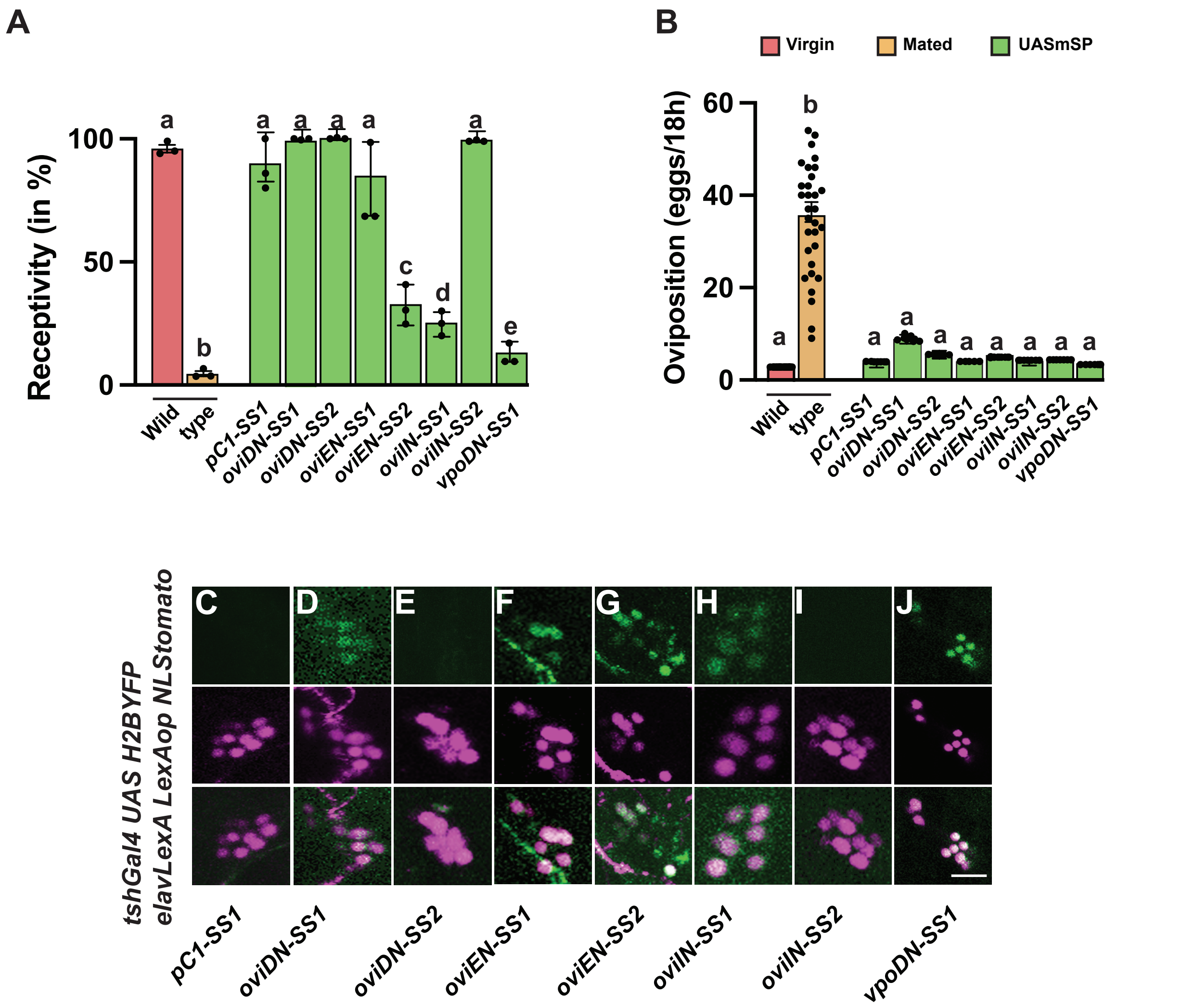
A
Receptivity (in %)
100
50
0
a
b
a
a
a
a
c
d
a
e
Wild
type
pC1-SS1
oviDN-SS1
oviDN-SS2
oviEN-SS1
oviEN-SS2
oviIN-SS1
oviIN-SS2
vpoDN-SS1
B
Virgin
Mated
UASmSP
Oviposition (eggs/18h)
60
40
20
0
a
b
a
a
a
a
a
a
a
a
Wild
type
pC1-SS1
oviDN-SS1
oviDN-SS2
oviEN-SS1
oviEN-SS2
oviIN-SS1
oviIN-SS2
vpoDN-SS1
tshGal4 UAS H2BYFP
elavLexA LexAop NLStomato
C
D
E
F
G
H
I
J
pC1-SS1
oviDN-SS1
oviDN-SS2
oviEN-SS1
oviEN-SS2
oviIN-SS1
oviIN-SS2
vpoDN-SS1

### Supp. Fig. S10

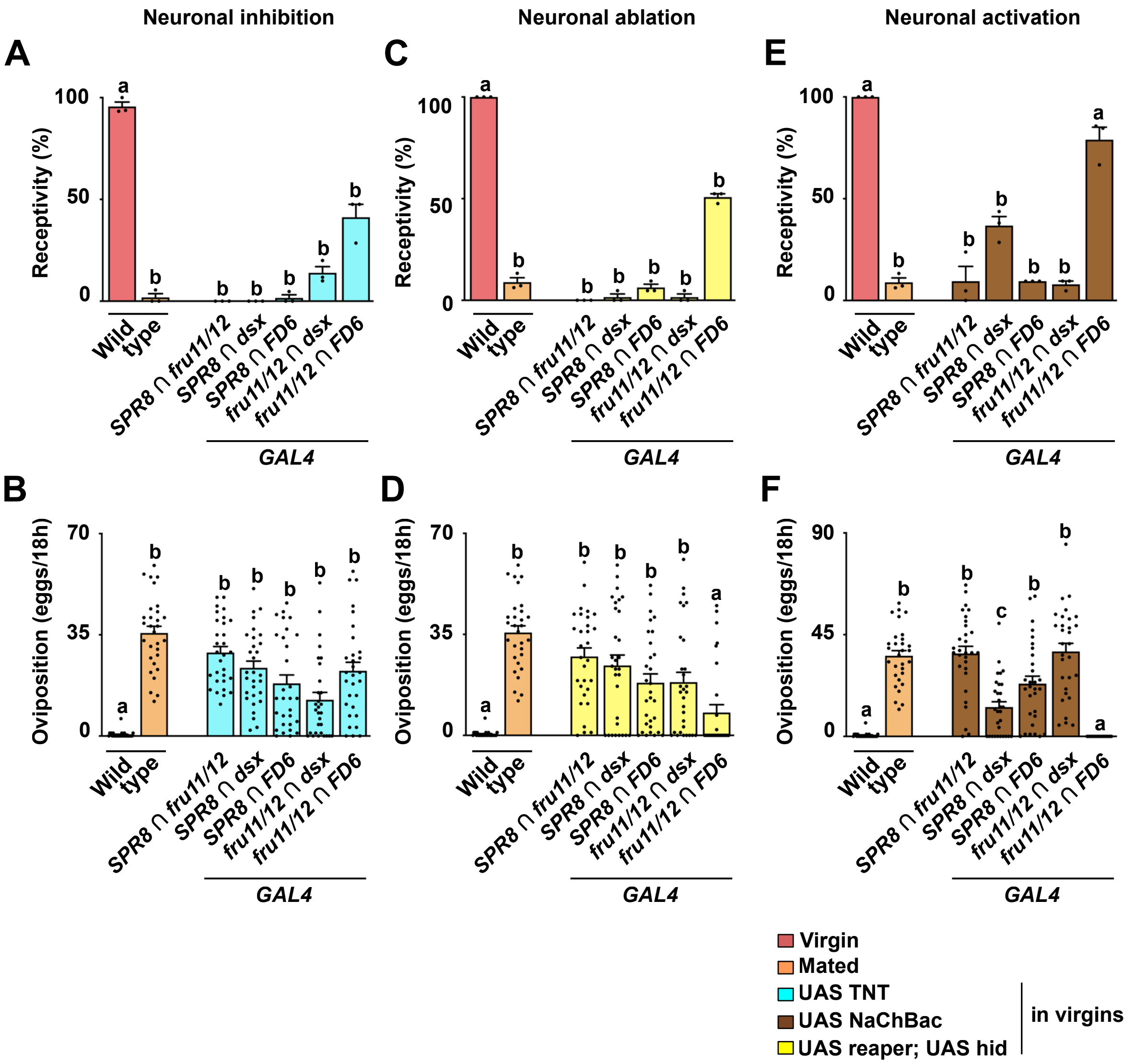

Neuronal inhibition
Neuronal ablation
Neuronal activation
A
B
C
D
E
F
Receptivity (%)
Oviposition (eggs/18h)
100
50
0
70
35
90
45
a
b
c
Wild type
SPR8 ∩ fru11/12
SPR8 ∩ dsx
SPR8 ∩ FD6
fru11/12 ∩ dsx
fru11/12 ∩ FD6
GAL4
Virgin
Mated
UAS TNT
UAS NaChBac
UAS reaper; UAS hid
in virgins

### Supp. Fig. S11

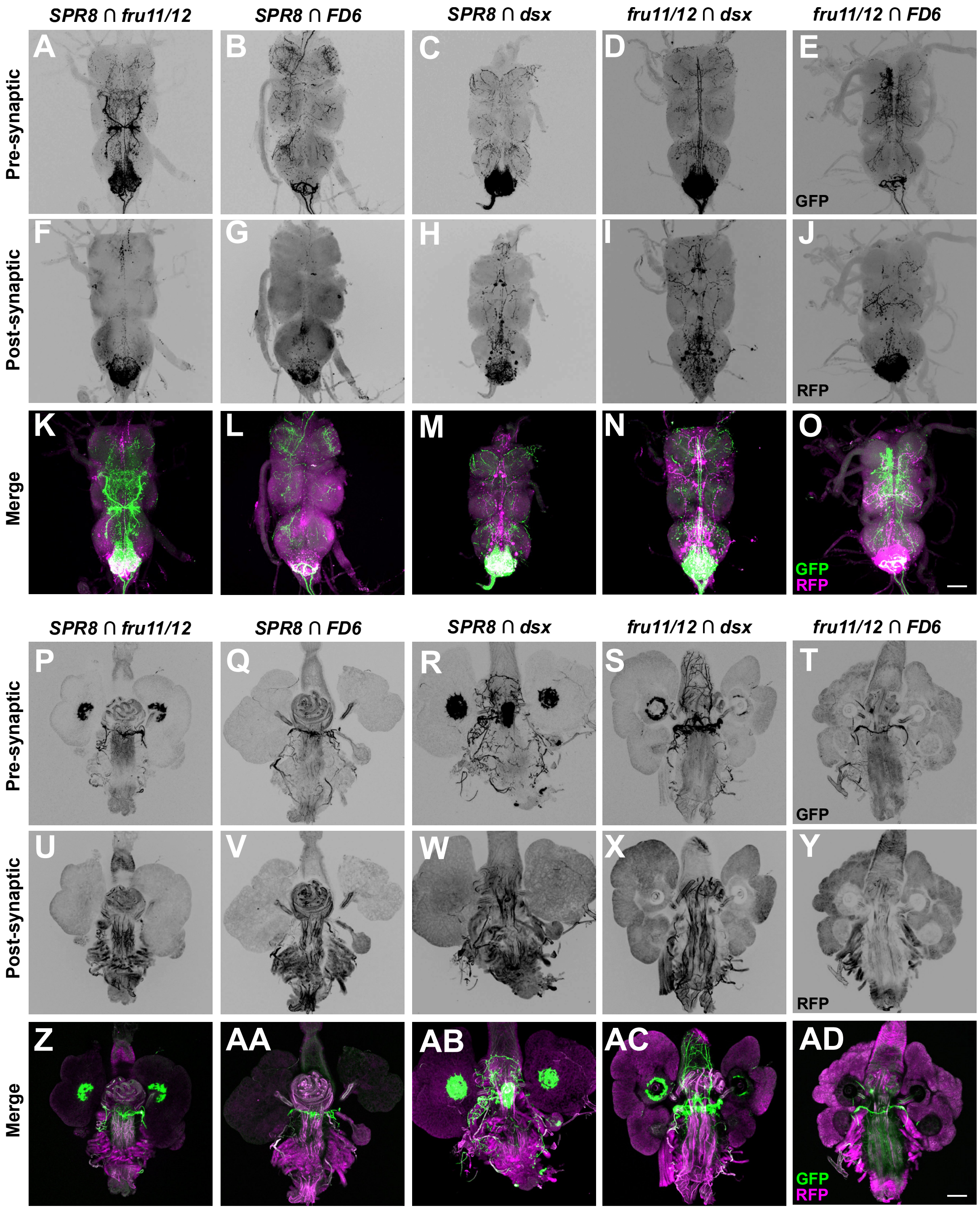

SPR8 ∩ fru11/12
SPR8 ∩ FD6
SPR8 ∩ dsx
fru11/12 ∩ dsx
fru11/12 ∩ FD6
Pre-synaptic
Post-synaptic
Merge
A
B
C
D
E
F
G
H
I
J
K
L
M
N
O
GFP
RFP
SPR8 ∩ fru11/12
SPR8 ∩ FD6
SPR8 ∩ dsx
fru11/12 ∩ dsx
fru11/12 ∩ FD6
Pre-synaptic
Post-synaptic
Merge
P
Q
R
S
T
U
V
W
X
Y
Z
AA
AB
AC
AD
GFP
RFP
